# Adipose-Derived Extracellular Vesicles Mitigate Experimental Cutaneous Leishmaniasis Through an IL-10–Dependent Mechanism

**DOI:** 10.64898/2026.02.02.703256

**Authors:** Douglas Barroso de Almeida, Camila Nascimento Fernandez, Alisson Amaral da Rocha, Igor Bittencourt dos Santos, João Victor Paiva Romano, Hozany Praxedes dos Santos, Naiara Carla dos Santos Manhães, Renata Trabach Santos, Beatriz Toja de Miranda, Luciana Covre, Alessandra Marcia da Fonseca Martins, Daniel Claudio Oliveira Gomes, Fernanda Ferreira Cruz, Tadeu Diniz Ramos, Patricia Rieken Macedo Rocco, Herbert Leonel de Matos Guedes

## Abstract

Leishmaniasis, a neglected tropical disease caused by protozoa of the *Leishmania* genus, is characterized by an imbalanced immune response that promotes parasite survival while inducing tissue damage. We previously showed that adipose-derived mesenchymal stem cells (AD-MSCs) limited lesion progression in C57BL/6 mice infected with *Leishmania amazonensis*. In the current study, we examined whether extracellular vesicles released by AD-MSCs (AD-MSC-EVs) reproduce these therapeutic effects. AD-MSC-EVs were purified, expanded, and characterized, confirming their typical morphology and EV-associated surface markers. *In vitro*, treatment of infected macrophages with AD-MSC-EVs 4 hours post-infection reduced parasite load through the induction of reactive oxygen species (ROS), independent of nitric oxide (NO). However, treatment 24 hours post-infection failed to reproduce this effect. *In vivo*, intralesional administration of AD-MSC-EVs significantly reduced lesion size without altering parasite load, while also decreasing levels of specific IgM and IgG for *Leishmania amazonensis* antigens. AD-MSC-EV therapy also reduced proinflammatory cytokines production by CD4⁺ and cytotoxicity CD8⁺ T cells, while increasing IL-10 production by γδ T cells. Moreover, combined therapy with pentavalent antimonial further reduced lesion size but did not affect parasite load. Notably, AD-MSC-EVs failed to control lesions in IL-10–deficient mice, suggesting that their therapeutic effect is dependent on IL-10. In conclusion, these findings showed that AD-MSC-EVs modulated the host immune response to attenuate *L. amazonensis*–induced cutaneous pathology and may represent a promising adjunct or alternative therapy for cutaneous leishmaniasis.

**Significance Statement:** Cutaneous leishmaniasis is a neglected tropical disease with limited therapeutic options and significant immunopathology. This study demonstrates that extracellular vesicles derived from adipose-derived mesenchymal stem cells (AD-MSC-EVs) can modulate host immune responses and reduce lesion severity in *Leishmania amazonensis* infection. Rather than directly eliminating parasites, AD-MSC-EVs attenuate tissue damage through IL-10 dependent immunoregulation, highlighting a novel host-directed therapeutic strategy. These findings advance understanding of EV-mediated immune modulation and support AD-MSC-EVs as a promising adjunct or alternative approach for treating cutaneous leishmaniasis and potentially other inflammatory infectious diseases.

## Introduction

Leishmaniasis is a group of neglected tropical diseases (NTDs) that threaten approximately 1 billion people worldwide and remain endemic in 99 countries across three continents (WHO, 2025). Infection with protozoa of the *Leishmania* genus produces a broad clinical spectrum, from self-limited cutaneous lesions to destructive mucocutaneous disease or life-threatening visceral involvement. In Brazil, *Leishmania (Leishmania) amazonensis*, a member of the *L. mexicana* complex, is an important etiologic agent of cutaneous leishmaniasis, responsible for localized and diffuse forms marked by chronicity and difficult therapeutic management (Rodríguez-Serrato MA, 2020). Disease severity reflects a dynamic interplay between parasite virulence factors and host immunity. *Leishmania* exploits dysregulated inflammatory responses and counter-regulatory cytokines to establish persistence, underscoring the immunopathological nature of the infection (Serrano-Coll H et al., 2025).

Despite expanded understanding of *Leishmania* immunobiology, current treatment options remain inadequate (Fischer T. et al., 2024). Infections caused by *L. amazonensis* are frequently refractory to standard therapy and associated with significant tissue damage. Available drugs are limited by toxicity, cost, emerging resistance (Bharadava K. et al., 2024), and the inability to correct harmful immune responses that support parasite survival. These limitations highlight the need for therapeutic strategies that combine antiparasitic activity with targeted immune modulation.

Mesenchymal stem cells (MSCs) have attracted interest as immunoregulatory therapies for inflammatory and infectious diseases (Wei S. et al., 2024; Habiba UE et al., 2024). Benefits have been reported in models of malaria (Thakur et al., 2020), Chagas disease (Mello DB et al., 2015), and leishmaniasis (Bahrami S et al., 2021). Our group previously showed that bone marrow–derived MSCs increased parasite load when administered intravenously in BALB/c mice, while intralesional delivery was ineffective (Pereira JC et al., 2017). More recently, we demonstrated that adipose-derived MSCs (AD-MSCs) reduced lesion progression in *L. amazonensis*–infected C57BL/6 mice and that combination with meglumine antimoniate further limited lesion expansion and parasite dissemination (Ramos et al., 2020). These findings suggest that MSCs act predominantly by modulating detrimental inflammation rather than exerting direct parasiticidal effects.

Concerns related to cell-based therapy: safety, biodistribution, differentiation potential, and standardization, have shifted attention to MSC-derived extracellular vesicles (EVs) (Regmi S. et al., 2019; Harrell et al., 2019; Pinheiro AAS et al., 2024). EVs are membrane-bound, non-replicating structures enriched with proteins, lipids, mRNAs, and microRNAs that mediate many reparative and immunomodulatory functions of MSCs (Song Y. et al., 2025; Quan J. et al., 2025). As a cell-free product, they circumvent risks associated with ectopic differentiation, embolization, and immune rejection (Kusuma et al., 2018).

A hallmark of *L. amazonensis* infection is profound dysregulation of cytokine networks with an induction of mixed Th1/Th2 (Ji et al., 2002), that limits effector T-cell responses, suppresses of macrophage activation,, and promotes parasite persistence (Maspi N, Abdoli A, Ghaffarifar F., 2016). Given that MSC-EVs influence macrophage phenotype and cytokine production, including IL-10–associated pathways, EVs derived from AD-MSCs represent a biologically plausible approach to restoring immune balance in cutaneous leishmaniasis.

Building on our observations that AD-MSCs improve lesion outcomes through immunomodulation, it remains essential to determine whether AD-MSC-derived EVs reproduce these effects and whether IL-10 is required for their activity. The present study therefore aimed to isolate, characterize, and evaluate AD-MSC-EVs in *L. amazonensis* infection, and to dissect their cytokine dependence using IL-10–deficient mice. By doing so, we sought to define the therapeutic relevance of AD-MSC-EVs and explore their potential as a safer, mechanistically defined, cell-free strategy for cutaneous leishmaniasis.

## Material and Methods

### Parasites

The parasites used in this study were *Leishmania (Leishmania) amazonensis* (MHOM/BR/75/Josefa strain), obtained from amastigotes isolated from lesions of infected BALB/c mice. The promastigotes were maintained at 26°C in M199 medium (Difco) supplemented with 10% fetal bovine serum (FBS; Cultilab) and cultured until the fifth passage for infection experiments. Infections were performed using stationary -phase promastigotes between the fourth and fifth days of culture.

### Preparation of MSCs

To extract mesenchymal stem cells (MSCs), male C57BL/6 mice (20–25 g, 8 weeks old) were used as donors. The animals were anesthetized with intravenous ketamine (25 mg/kg; Sigma) and xylazine (2 mg/kg; Sigma) and euthanized by cervical dislocation. Using sterile forceps and scissors, an abdominal incision was made to remove the testicles. The epididymal fat surrounding the testis was extracted and suspended in phosphate-buffered saline (PBS).

To isolate adipose-derived MSCs (AD-MSCs), epididymal fat pads were collected, rinsed with PBS, transferred to a Petri dish, and cut into small fragments (0.2–0.8 cm²). The tissue was further minced, washed with PBS, and enzymatically digested with type I collagenase (1 mg/mL in PBS) for 30–40 min at 37°C. After digestion, the suspension was allowed to settle for 1–3 min to remove undigested tissue. The supernatant was transferred to a new tube, mixed with fresh medium, and centrifuged at 400×g for 10 min at 25°C. The resulting pellet was resuspended in 1 mL of Iscove’s Modified Dulbecco’s Medium (IMDM; Invitrogen) containing 1% penicillin-streptomycin (5000 IU/mL and 5000 μg/mL, respectively; Gibco) and 20% heat-inactivated FBS (Invitrogen). Cells were seeded in T25 flasks (4 mL per flask) and incubated at 37°C in a humidified atmosphere with 5% CO₂. On day 3, the medium was changed to remove non-adherent cells. Once adherent cells reached 80% confluence, they were passaged using a 0.25% trypsin-EDTA solution (Gibco) and maintained in complete IMDM medium (IMDM with 20% FBS and 1% penicillin-streptomycin).

### Isolation of Extracellular Vesicles from MSCs

MSCs were expanded in culture until passage five. Once cells reached 70–90% confluence, they were maintained in serum-free IMDM for 48 hours. The conditioned medium was collected and processed as follows: Centrifugation at 400×g for 10 min to remove large cells and debris, Centrifugation at 3000×g for 20 min to remove larger particles (>1000 nm), Filtration through a 0.22-μm membrane to eliminate residual contaminants, and Ultracentrifugation at 100,000×g for 2 hours (modified from Welsh JA et al., 2024) to isolate extracellular vesicles (EVs). The EV pellet was washed, resuspended in PBS, and stored at - 80°C for future use.

### Characterization of Extracellular Vesicles

EVs were characterized using three methodologies recommended by the International Society for Extracellular Vesicles (ISEV) (Welsh JA *et al*., 2024).

1. Flow Cytometry: Fifty microliters of a mix containing EV samples from three differents isolations were incubated with 0.25 μL of latex beads (4 μm diameter, 5.5 × 10^9^ particles/mL) for 20 min at room temperature. The sample was then incubated overnight in 1 mL of FACS Buffer (PBS with 0.1% BSA and 0.01% NaN₃) with constant rotation. After washing, EV-bound beads were stained with antibodies against CD9^+^ APC (BioLegend #312107), CD63^+^ PE (BioLegend #353004), and CD81^+^ PE (BioLegend #104905) for 45 min at 4°C.Stained samples were analyzed using a BD FACS Canto II flow cytometer.
2. Nanoparticle Tracking Analysis (NTA): Samples were diluted 1:50 in sterile PBS (1 mL final volume) and analyzed using a NanoSight NS300 system (Malvern Panalytical). Five 60-second videos were captured under standardized settings: temperature 25°C, syringe speed 25 μL/s, camera level 13, screen gain 1.0. Data were analyzed using NTA software (NTA 3.4 buildt 3.4.4).
3. Transmission Electron Microscopy (TEM): EV samples were placed on carbon-coated formvar grids, stained with 5% uranyl acetate, and examined using a JEOL 1200 EX electron microscope (Jeol LTD).

### Peritoneal Macrophage Culture

C57BL/6 mice were anesthetized with isoflurane and euthanized by cervical dislocation. The peritoneal cavity was washed with 5 mL of cold RPMI-1640 (Sigma) using a 24G needle and syringe. The lavage fluid was collected, kept on ice, and centrifuged at 400×g for 5 min at 4°C. The pellet was resuspended in 5 mL of RPMI-1640 without FBS, and total cells were counted using a Neubauer chamber. Depending on the macrophage yield, either 5×10^5^ or 1×10^6^ cells were plated per well in 24-well plates and incubated at 37°C with 5% CO₂ for 1 hour to allow cell adhesion. Non-adherent cells were removed by washing three times with PBS at 37°C, and adherent cells were cultured in complete RPMI medium (1% glutamine, 1% essential amino acids, 1% pyruvate, 1% β-mercaptoethanol, and 10% FBS) for 24 hours.

### In vitro infection, treatment, and ROS/NO quantification

After 24 hours of the previous procedure, the plated macrophages were washed three times with PBS at 37°C, and 350 µL of complete RPMI medium was added to the wells. The macrophages were then infected with *L. amazonensis* at a ratio of 5 or 10 parasites per macrophage for 4 hours. After this incubation period, the wells were washed three times with PBS at 37°C, and 600 µL of complete RPMI medium was added. Next, 50 µL of PBS or extracellular vesicles were added to the cells in the following amounts: 1×10^6^, 5×10^6^, 1×10^7^, 1×10^8^, or 1×10^9^ per well. The macrophages were then incubated at 37°C with 5% CO_2_ for 24, 48, or 72 hours. After the incubation period, the supernatant was collected for nitric oxide (NO) measurement, and the cells were stained using the rapid Panoptic kit (LaborClin) and manually counted in groups of 100 cells per well slide using an optical microscope (Olympus, CX31) at 100x magnification. The following parameters were calculated:

- Percentage of infected cells (% macr): This was obtained by counting the number of infected macrophages out of the 100 macrophages counted.
- Total number of amastigotes (parasite load): This was calculated by counting the total number of amastigotes present in all macrophages.

NO measurement was performed using the Griess Reagent method, using nitrite as a standard curve with concentrations of 50, 25, 12.5, 6.25, 3.125, 1.5625, and 0.78125 µM. ROS production levels were assessed using the Dihydrorhodamine 123 (DHR 123) probe and detected by spectroscopy.

### *In vivo* infection and lesion monitoring

Animals were infected intradermally with 10 µL of PBS containing 2×10^6^ *L. amazonensis* promastigotes in the footpad of the right hind paw. Lesion growth was monitored by measuring the thickness of the infected paw (in mm) weekly with a thickness gauge (Mitutoyo), discounting the initial measurement before infection.

### Treatment and administration of extracellular vesicles derived from AD-MSCs

The mice received three doses of 1×10^9^ extracellular vesicles derived from adipose mesenchymal stem cells (AD-MSCs) for the treatment of infection via a retro-orbital intravenous or intralesional route. The animals were anesthetized with isoflurane and received PBS or EVs through the orbital plexus or in the infected paw. The first dose was administered 14 days after infection, the second dose on day 21, and the third dose on day 28 post-infection.

### Administration of meglumine antimoniate

Treatment was initiated on the 18th day post-infection. On alternate days (Mondays, Wednesdays and Fridays) the animals received 2 mg of meglumine antimoniate (100 mg/kg) in 100 μL of PBS at a concentration of 10 mg/mL. The drug was injected intraperitoneally.

### Specific antibody production

Quantification of antibody production was performed using ELISA (Goat Anti-Mouse IgM-UNLB:Cat. No. 1021-01, Goat Anti-Mouse IgG Fc-UNLB:Cat. No. 1033–01, Goat Anti-Mouse). Total *L. amazonensis* antigen (LaAg) diluted in PBS (5 μg/mL) was added to the plate for coating overnight. On the second day, the content was discarded, and the plate was blocked with Block Buffer (PBS with 5% heat-inactivated fetal bovine serum [HIFCS, GIBCO Laboratories, Grand Island, NY, USA] and 0.05% Tween 20) for 1 hour. The plate was then washed three times with Wash Buffer (PBS with 0.05% Tween 20), and the samples were diluted in Block Buffer and added to the plate. After 1 hour, the plate was washed five times with Wash Buffer, and the secondary antibody specific for each isotype was added. After 1 hour, the plate was washed seven times with Wash Buffer, and TMB was added. The reaction was blocked with HCl.

### Detection of cytokines by enzyme-linked immunosorbent assay (ELISA)

Concentrations of the cytokines present in cell culture supernatants and the footpad homogenate supernatants were determined by the ELISA using commercial ki7s (Preprotech) according to the datasheet instructions. Recombinant murine IL-4, IL-10, IL-10, IFN-γ, and IL-22 were used to generate the standard curves.

### Determination of parasite load

The mice were anesthetized with isoflurane and euthanized by cervical dislocation. The infected paw of each animal was removed and placed in a 70% alcohol bath for 1 minute before being individually weighed. The spleens and lymph nodes were then removed and placed in individual Eppendorf tubes containing 1 mL of M199 (Difco) and RPMI-1640 (Sigma) medium, respectively. The number of parasites in the infected paw and organs of the animals was determined using the limiting dilution technique. Paw and organ homogenates were obtained by manual maceration, adding 1 mL of M199 medium (Difco) for the paws and the respective media for the other organs. Subsequently, 50 µL of sample homogenates were serially diluted in duplicate in 96-well culture plates, containing 150 µL of M199 (Difco) 10% SFB (Cultilab) in each well (1:4 dilution per well), and incubated at 26°C for 15 days to monitor promastigote growth. The cultures were observed under an optical microscope on days 7 and 14 after plating, and the last dilution showing promastigotes (limiting dilution) was recorded. The total parasite load was calculated using the formula: (4^x^)/g, where x = number of the last well where promastigotes were observed and g = footpad or organ weight.

### Cell staining for flow cytometry

Draining lymph nodes from the lesion (popliteal) were individually removed and manually macerated. The cytokine production of lymph node cells was evaluated after polyclonal activation, as *LaAg* induces apoptosis of T cells (Pinheiro et al., 2004). Cells (1 × 10^6^/well) in a 96-well plate were stimulated for 4 hours at 37°C with PMA (phorbol 12-myristate 13-acetate, 10 ng/mL, Sigma-Aldrich) and Ionomycin (10 ng/mL, Sigma-Aldrich). All centrifugation steps were performed at 4°C. Cells were washed with PBS and blocked with 50 µL/well Human FcX (BioLegend) for 15 minutes. Then, 50 µL/well of the staining antibody pool was added and incubated for 30 minutes at 4°C. Cells were washed with buffer solution (PBS with 5% FBS) at 400 g for 5 minutes, then fixed and permeabilized using the FoxP3 permeabilization/fixation kit (eBioscience) according to the manufacturer’s protocol. After another wash, cells were resuspended in the same solution. For extracellular staining, the following antibodies were added (all used at 0.1 μg/ml) and incubated for 1 hour at 4°C in the dark: CD3 (anti-CD3-BV-421; clone 17A2, BioLegend), CD4 (anti-CD4-BV-785; clone RM4-5, BioLegend), CD8 (anti-CD8-PerCP/Cyanine 5.5; clone 53-6.7, BioLegend), CD107a (anti-CD107a-PE/Cyanine 7; clone 1D4B, BioLegend). For intracellular staining, the following antibodies were added (all used at 0.1 μg/ml) and incubated for 1 hour at 4°C in the dark: Perforin (anti-Perforin-PE; clone S16004A, BioLegend), Granzyme (anti-Granzyme-Alexa Fluor 647; clone GB11, BioLegend), IFN-γ (anti-IFN-γ-APC; clone XMG1.2, eBioscience), IL-17 (anti-IL17-APC/Cyanine 7; clone TC11-18H10.1, BioLegend), and IL-10 (anti-IL-10-PE; clone JES5-16E3, BioLegend). After washing, cells were resuspended in 200 μL buffer solution and stored in the dark at 4°C until acquisition. Cells were analyzed on a BD LSR Fortessa flow cytometer; data from 100,000 events were captured from cells acquired on CD3+ and analyzed with FlowJo® software (BD-Becton, Dickinson & Company).

### Statistical Analysis

Sample size was based on pilot studies. Subsequently, pilot studies were performed using the current model to confirm the sample size adequacy for assessing footpad damage. This approach aligns with the 3Rs principle, ensuring that the use of animals was minimized while maintaining the scientific integrity of the study. Student t-test was used to compare 2 groups, when more groups were compared, One way ANOVA was used. Data that satisfied parametric assumptions were expressed as the mean and standard deviation and data that did not satisfy parametric assumptions as the median (interquartile range). All tests were carried out in GraphPad Prism 8 (GraphPad Software, La Jolla, CA, United States). Significance was established at *p* < 0.05.

## Results

### Production and characterization of AD-MSC–derived extracellular vesicles

AD-MSC–EVs were isolated and characterized prior to experimental use. Transmission electron microscopy showed vesicles with the expected spherical morphology and heterogeneous diameters (Figure 1A). Nanoparticle tracking analysis confirmed a heterogeneous size distribution ranging from approximately 30 to 500 nm, with a mean diameter of 195.7 nm and an average concentration of 1.1 × 10^10^ particles/mL (Figure 1B). EV surface markers were assessed by bead-based flow cytometry. Unlabeled controls displayed >99.9% double-negative staining, confirming low background (Figure 2A). CD9-, CD63-, and CD81-labeled samples demonstrated clear positive signal shifts relative to controls, with 95.7%, 96.7%, and 93.8% positivity, respectively (Figure 2B–G). These findings confirmed the expected identity and purity of the EV preparations.

**Figure 1.**
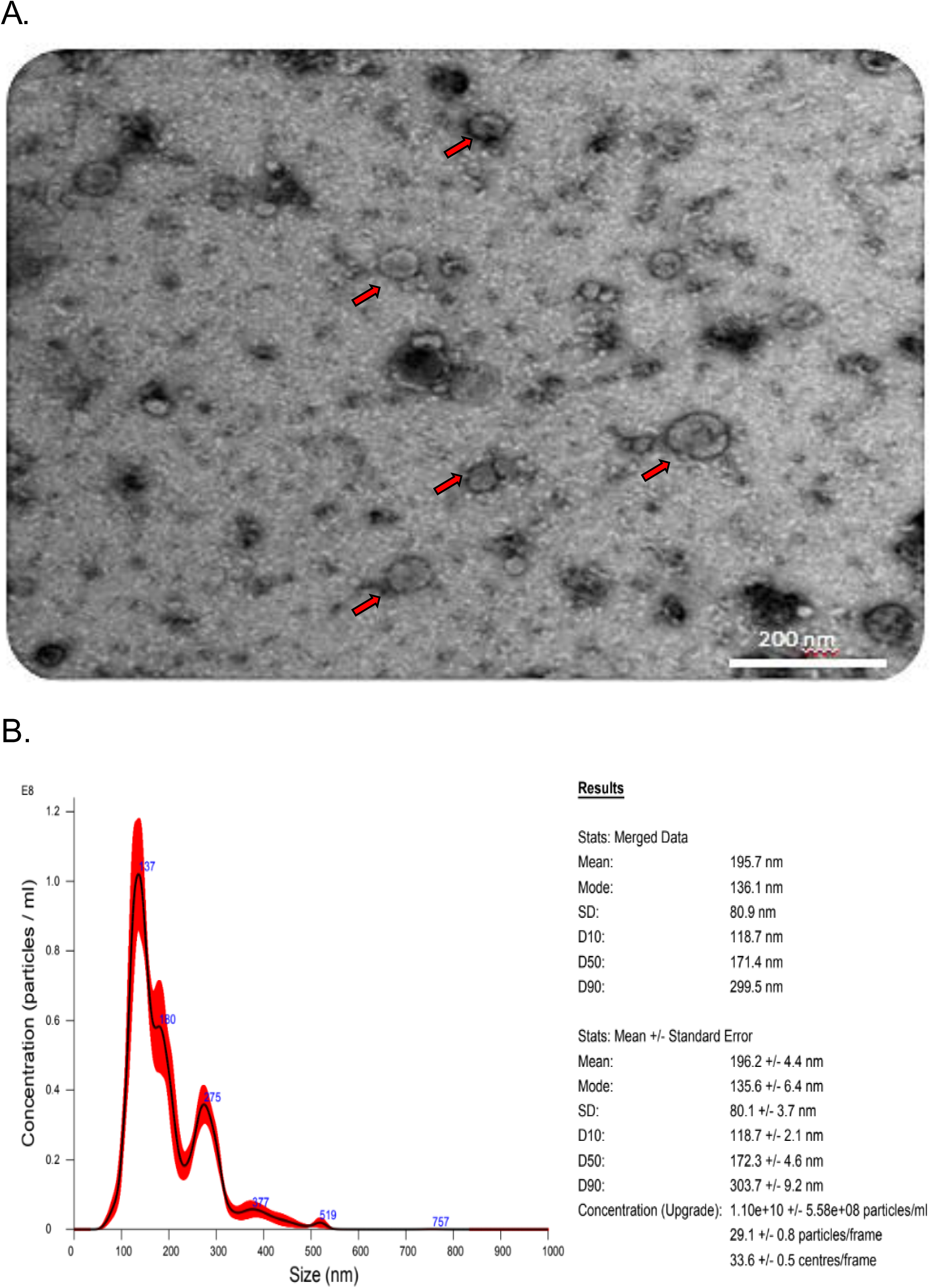
Characterization of adipose-derived MSC extracellular vesicles (EVs). (A) Representative transmission electron microscopy image of isolated EVs. (B) Representative nanoparticle tracking analysis profile of EV preparations. Red arrows indicate vesicular structures.

**Figure 2.**
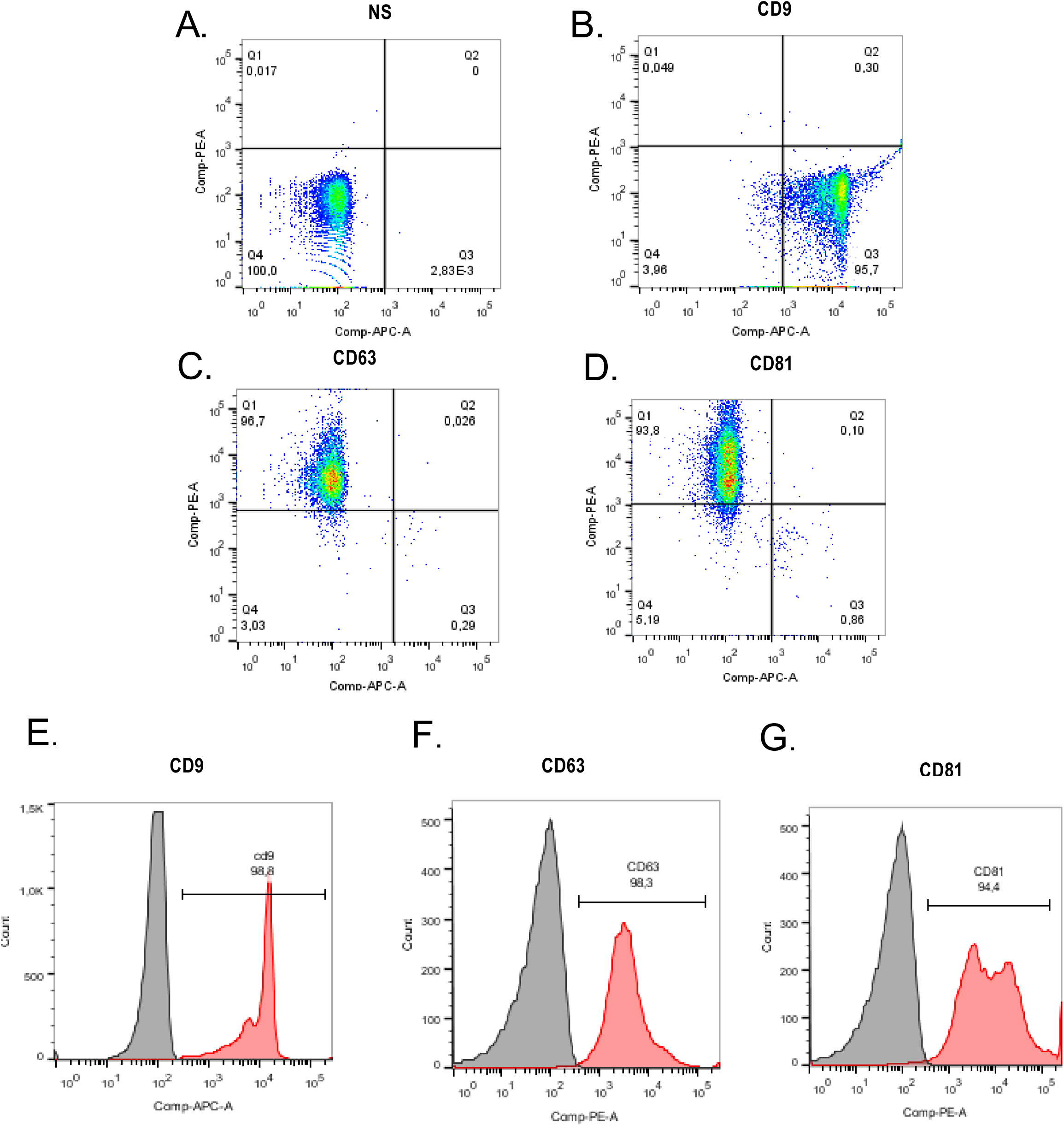
Flow cytometric profiling of adipose-derived MSC EVs. EVs from three independent isolations were pooled and stained with CD9 (APC), CD63 (PE), or CD81 (PE). (A) Unstained EVs; (B) CD9-stained EVs; (C) CD63-stained EVs; (D) CD81-stained EVs. (E–G) Histograms showing fluorescence intensity of CD9, CD63, and CD81 (red) compared with unlabeled beads (grey).

### AD-MSC-EVs reduce parasite load in infected macrophages in a time- and dose-dependent manner

To evaluate the early impact of EVs on *L. amazonensis* infection, macrophages were treated 4 h post-infection with escalating EV doses. Only the highest initial dose (1 × 10⁷ EVs/well) significantly reduced both the percentage of infected macrophages and total parasite load at 24 and 48 h, without affecting nitric oxide production (Figure S1). Subsequent experiments using 1 × 10⁷, 1 × 10⁸, and 1 × 10⁹ EVs/well confirmed a dose-dependent reduction in infection at 24, 48, and 72 h (Figure 3A–F). Parasite load decreased across all time points, whereas NO levels remained unchanged (Figure 3G–I). Treatment with 1 × 10⁹ EVs increased ROS production at 24 h (Figure 3J). To assess whether macrophage activation influenced EV efficacy, cells were pre-stimulated with IFN-γ (0.5 ng/mL). IFN-γ partially blunted the EV antiparasitic effect, which remained detectable only at 1 × 10⁹ EVs (Figure S2D–E). IFN-γ pre-stimulation increased NO production following EV exposure (Figure S2F). Finally, EV administration at 24 h post-infection failed to modify infection parameters or ROS generation, in contrast to treatment at 4 h, which reduced infection rate, decreased parasite load, and increased ROS (Figure 4A–F). These findings indicate that EV-mediated control of parasite load is restricted to early stages of infection.

**Figure 3.**
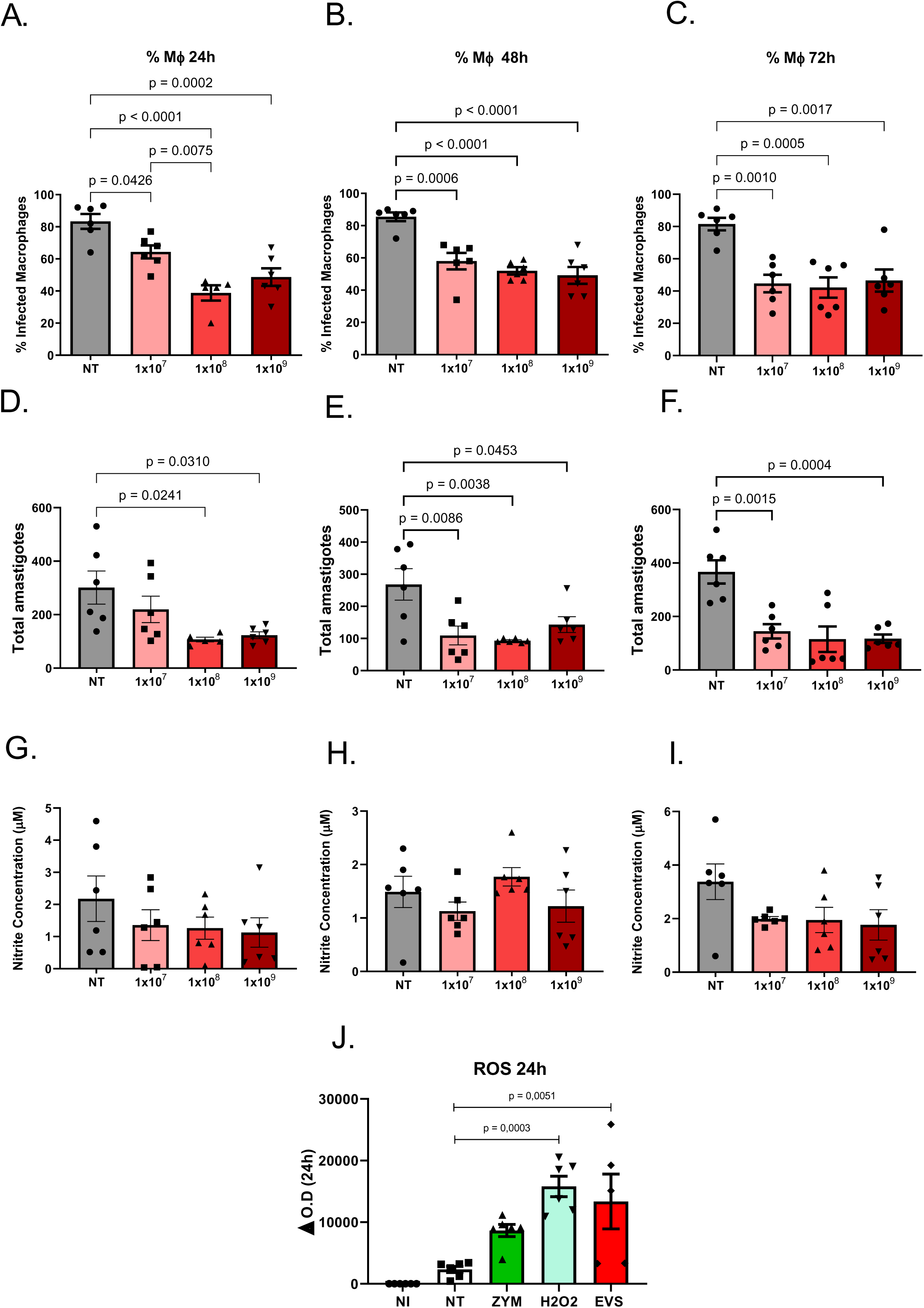
Effects of MSC EVs on peritoneal macrophages infected with *L. amazonensis*. Macrophages were infected for 4 h and then treated with EVs for 24, 48, or 72 h. (A, D, G) Percentage of infected macrophages; (B, E, H) Number of internalized amastigotes per 100 macrophages; (C, F, I) Nitric oxide levels in culture supernatants; (J) ROS levels at 24 h. Data are mean ± SEM from two or three independent experiments. NI: non-infected; NT: infected, untreated; ZYM: zymosan; EVs: 10⁹ EVs; FORM: formalin-fixed EVs. One-way ANOVA with Tukey’s post-test.

**Figure 4.**
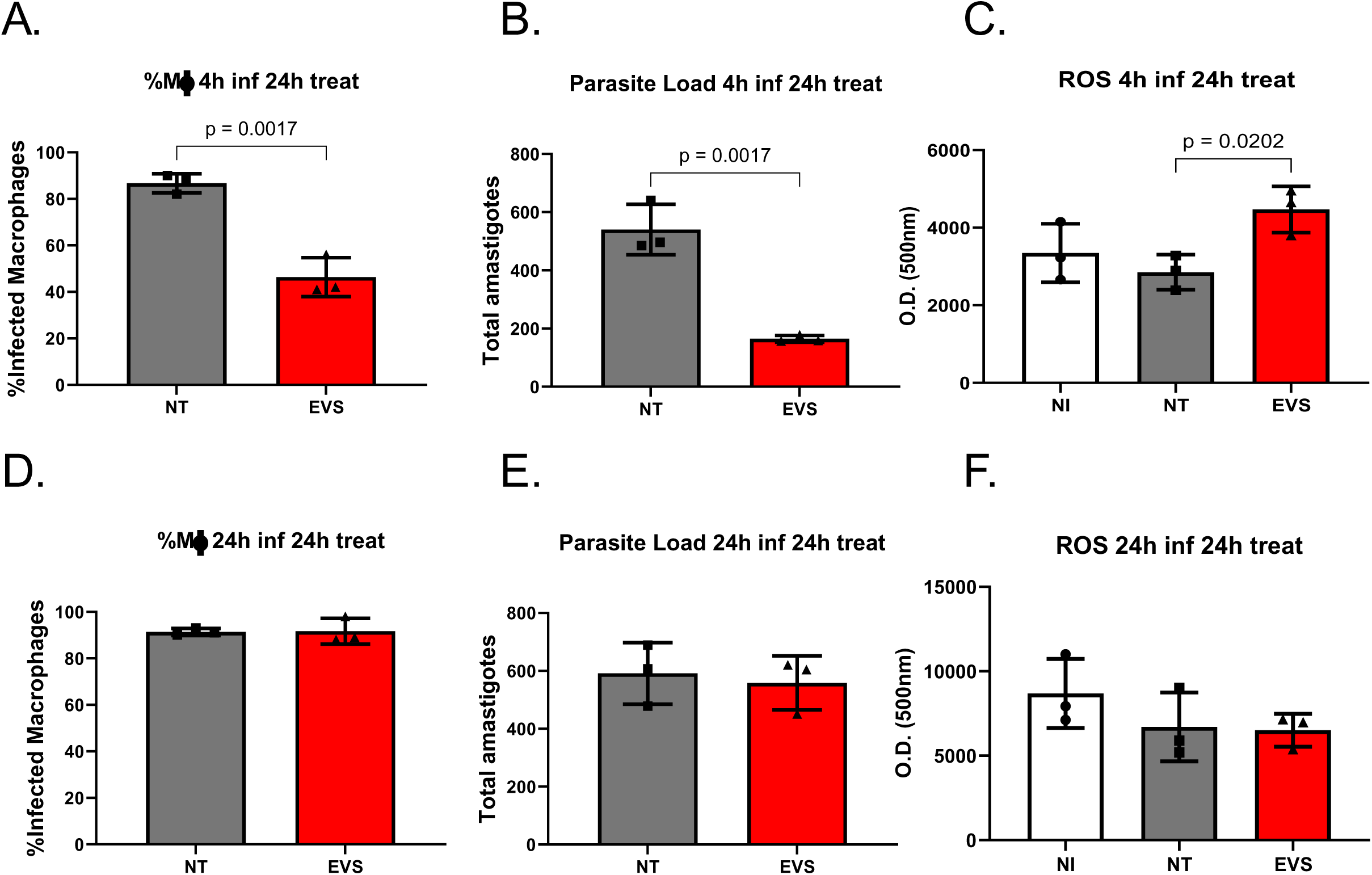
MSC EV treatment at early or late infection time points in peritoneal macrophages. Macrophages were infected for 4 h or 24 h and treated with 10⁸ EVs for 24 h. (A, D) Percentage of infected macrophages; (B, E) Internalized amastigotes per 100 macrophages; (C, F) ROS levels. Data are mean ± SEM from two independent experiments. NI: non-infected; NT: infected, untreated. Only p < 0.05 is shown. Unpaired t-test.

### Intralesional AD-MSC-EV treatment limits lesion progression but does not alter parasite load in C57BL/6 mice

To evaluate the *in vivo* efficacy of EV treatment, C57BL/6 mice infected in the hind footpad received EVs either intravenously or intralesionally (1 × 10⁹ EVs per dose) on days 14, 21, and 28 post-infection. Intravenous administration produced no measurable effect on lesion size compared with control animals (Figure 5A). In contrast, intralesional administration significantly reduced lesion progression beginning one week after the first treatment, with this effect maintained through the peak inflammatory phase (Figure 5A). Representative images from infected paws shows that the group who received intralesional treatment had smaller lesion with a milder inflammatory profile (Figure 5B). Thereafter, lesion trajectories converged among all groups. Despite the improvement in lesion size, parasite burdens in the infected footpad, spleen, and draining lymph nodes were not significantly different between treated and control animals (Figure 5C–E). Together, these findings indicate that EV treatment can limit lesion development without affecting parasite load.

**Figure 5.**
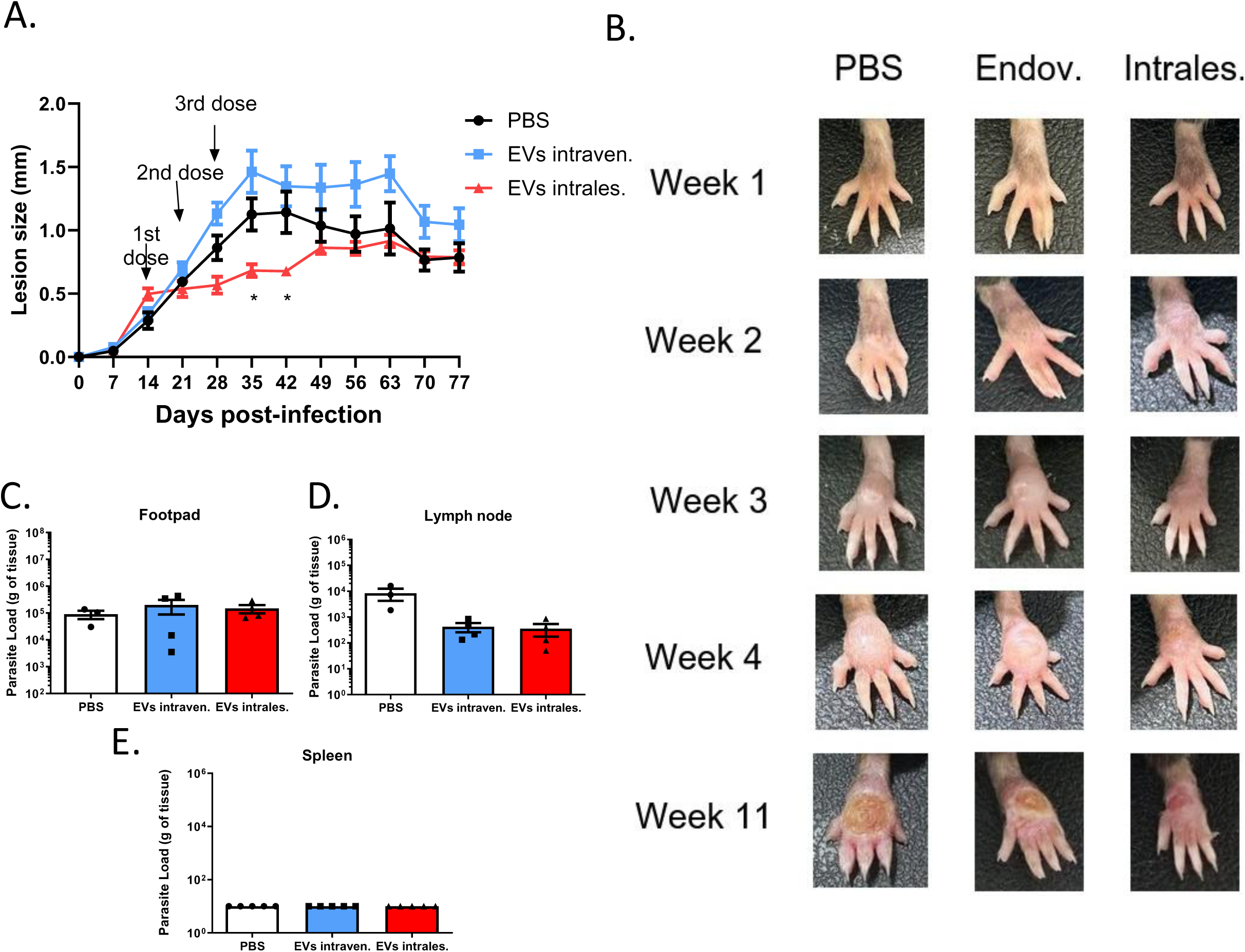
Systemic or intralesional administration of MSC EVs in *L. amazonensis*-infected mice. C57BL/6 mice were infected with 2×10⁶ promastigotes. EVs (10⁹) were administered intravenously or intralesionally on days 14, 21, and 28. (A) Lesion progression; (B) Representative paw images; (C–E) Parasite load in footpad, lymph nodes, and spleen at day 77. n=3 (PBS), n=4 (other groups). *p < 0.01, **p < 0.001, ***p < 0.0001 compared with PBS. Two-way ANOVA with Bonferroni post-test.

### AD-MSC-EVs modulate humoral responses and T cell cytokine profiles

To investigate the immunomodulation of EVs *in vivo*, we performed the treatment using the same protocol and euthanized at the peak infection. Intralesional EV treatment reduced lesion size (Figure 6A), a finding corroborated by representative footpad images showing smaller and less edematous lesions in treated mice (Figure 6B), however, parasite load remained unchanged in footpad, spleen, and lymph nodes (Figure 6C–E). EV-treated mice serum exhibited lower total IgG and IgM reactivity against *L. amazonensis* antigens compared with PBS controls (Figure 6G–H).

**Figure 6.**
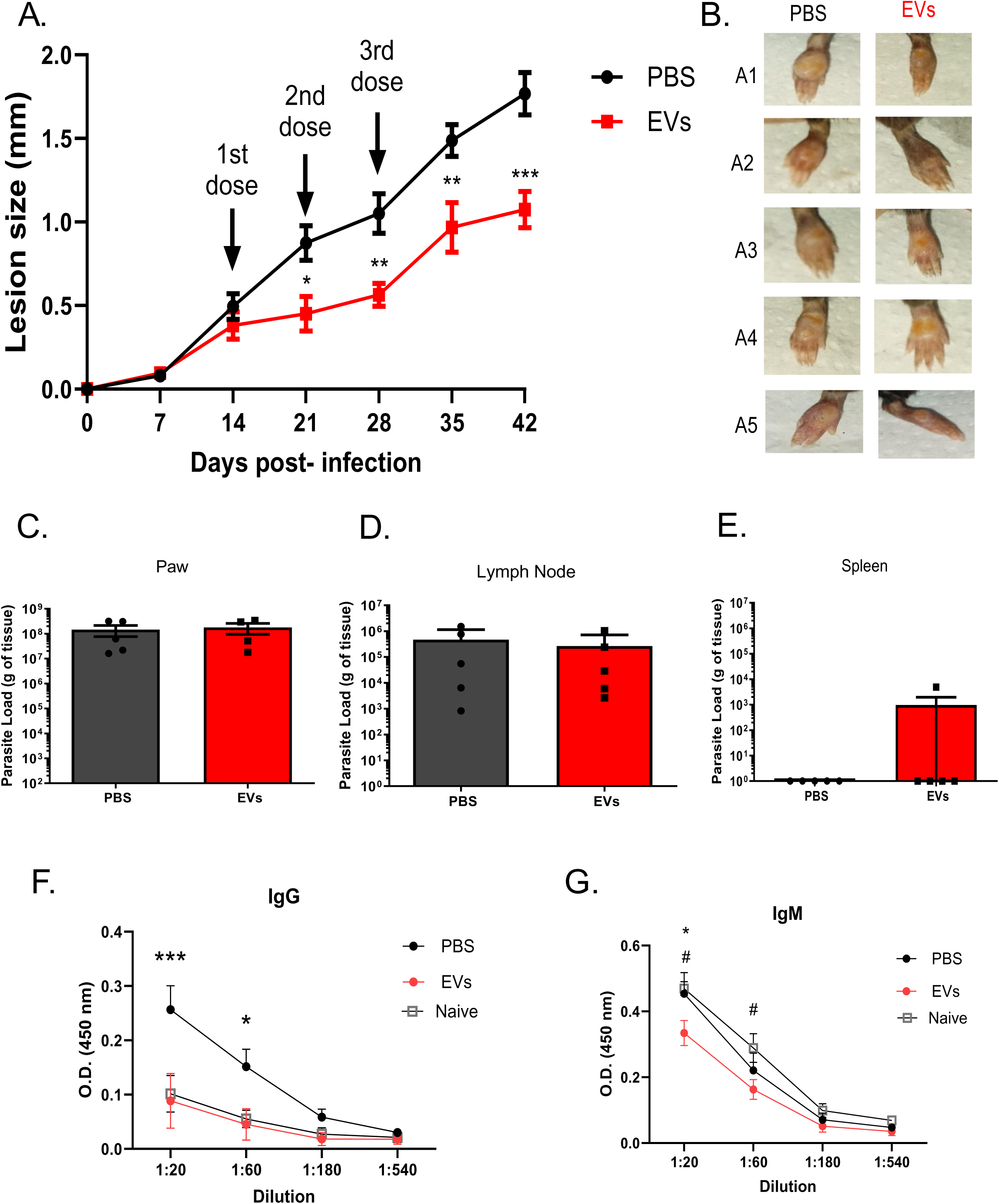
Intralesional MSC EV therapy in *L. amazonensis*-infected mice. EVs (10⁹) were administered intralesionally on days 14, 21, and 28. (A) Lesion progression; (B) Representative paw images; (C–E) Parasite load in footpad, lymph nodes, and spleen at day 43; (F–G) Serum total IgG and IgM. Data are mean ± SEM from one of three experiments. *p < 0.01, **p < 0.001, ***p < 0.0001 vs. PBS; #p < 0.01 vs. naïve. Two-way ANOVA with Bonferroni post-test.

Flow cytometric analysis of draining lymph node cells was done using the same gating Strategy for all analysis (Figure S3) and revealed reduced frequencies and absolute numbers of IFN-γ– and IL-17–producing CD4⁺ T cells following EV treatment (Figure 7C–F). IL-10 production by CD4⁺ T cells was not altered (Figure 7G–H). However, EV treatment increased the frequency of IL-10–producing CD3⁺ T cells lacking CD4 and CD8 expression (Figure 8A–B). Subsequent gating identified these cells as γδ T cells, which selectively upregulated IL-10 production (Figure 8C). Notably, EV treatment did not affect the frequency or absolute number of γδ T cells compared with PBS controls (Figure 8E–F), nor did it significantly modulate their production of other cytokines (Figure 8G–J). No changes were detected in cytokine production by CD8⁺ T cells (Figure S5).

**Figure 7.**
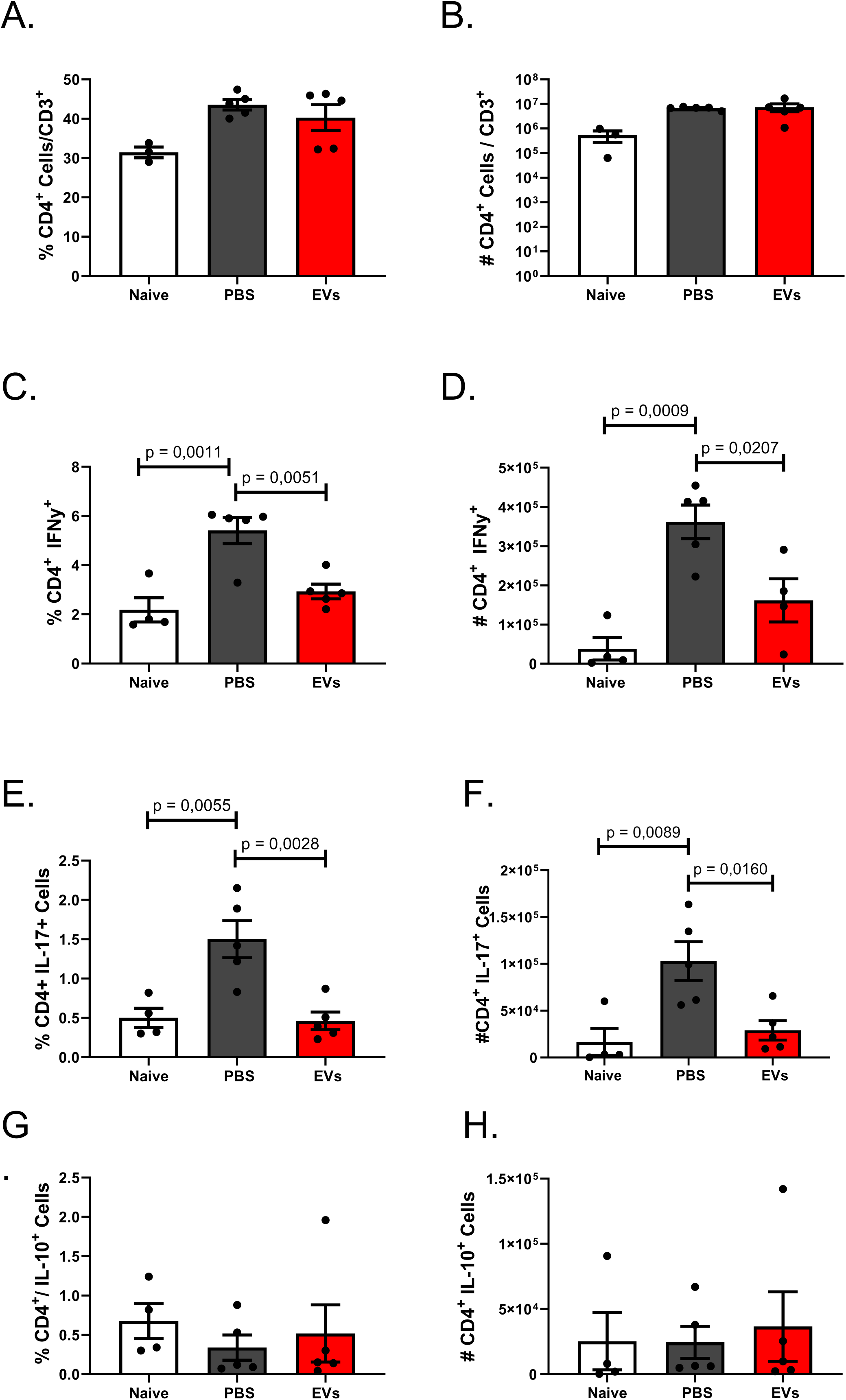
CD4⁺ T-cell responses following intralesional MSC EV therapy. Draining lymph nodes were harvested on day 43. (A, B) Frequency and number of CD4⁺ T cells; (C, D) IFN-γ⁺CD4⁺ cells; (E, F) IL-17⁺CD4⁺ cells; (G, H) IL-10⁺CD4⁺ cells. n=4 (naïve), n=5 (PBS, EVs). Only p < 0.05 is shown. One-way ANOVA with Tukey’s post-test.

**Figure 8.**
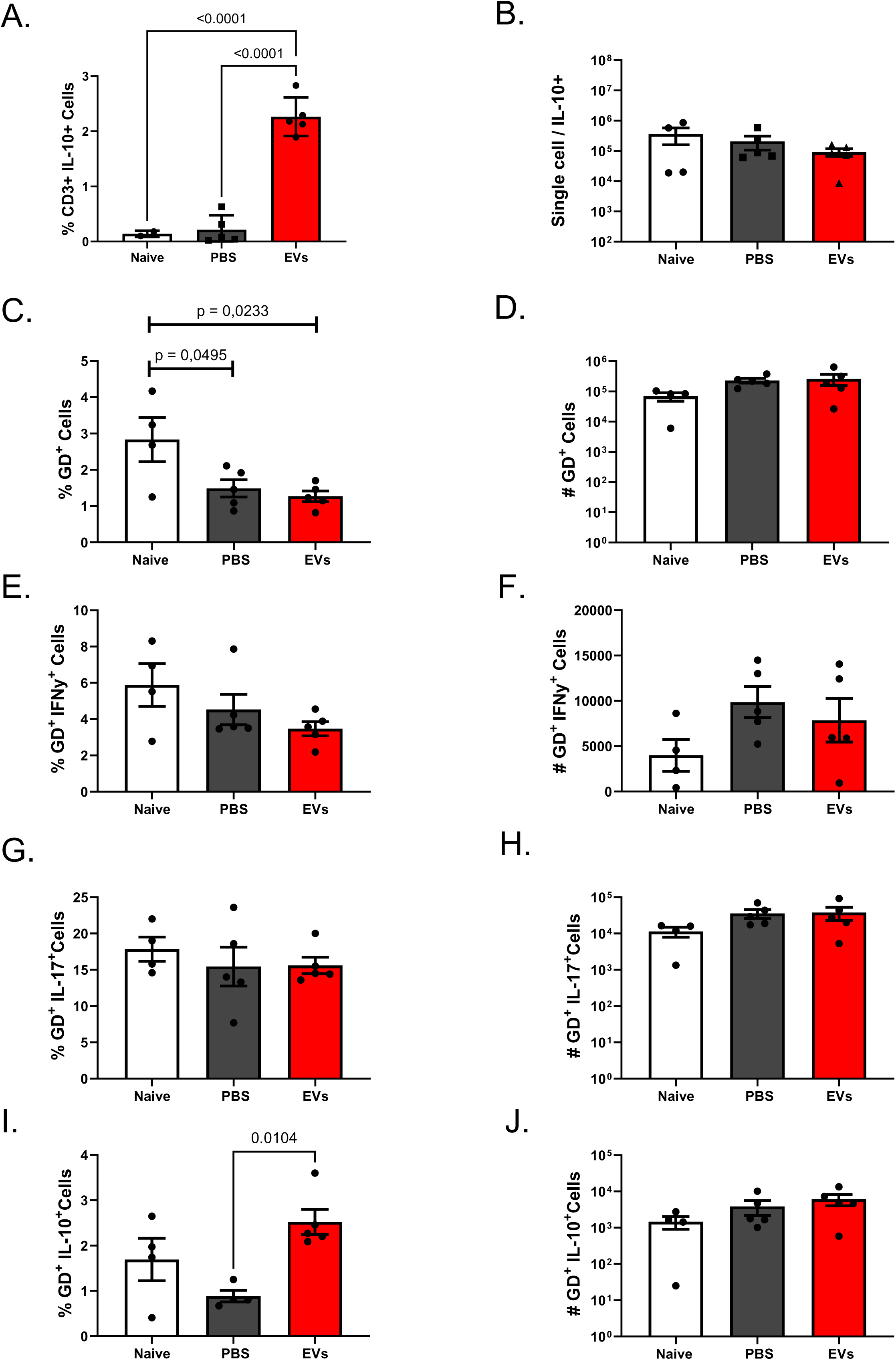
γδ T-cell cytokine responses following MSC EV therapy. Draining lymph nodes were collected on day 43. (A, B) IL-10⁺CD3⁺ cells; (C, D) IL-10⁺γδ T cells; (E, F) Frequency and number of γδ T cells; (G–J) IFN-γ⁺ and IL-17⁺ γδ T cells. n=4 (naïve), n=5 (PBS, EVs). Only p < 0.05 is shown. One-way ANOVA with Tukey’s post-test.

Regarding cytotoxic activity of CD8⁺ T cells, EV-treated mice showed reduced numbers of granzyme B–producing CD8⁺ T cells (Figure 9D) and decreased CD107a expression (Figure 9H), whereas perforin expression was unchanged (Figure 9E–F).

**Figure 9.**
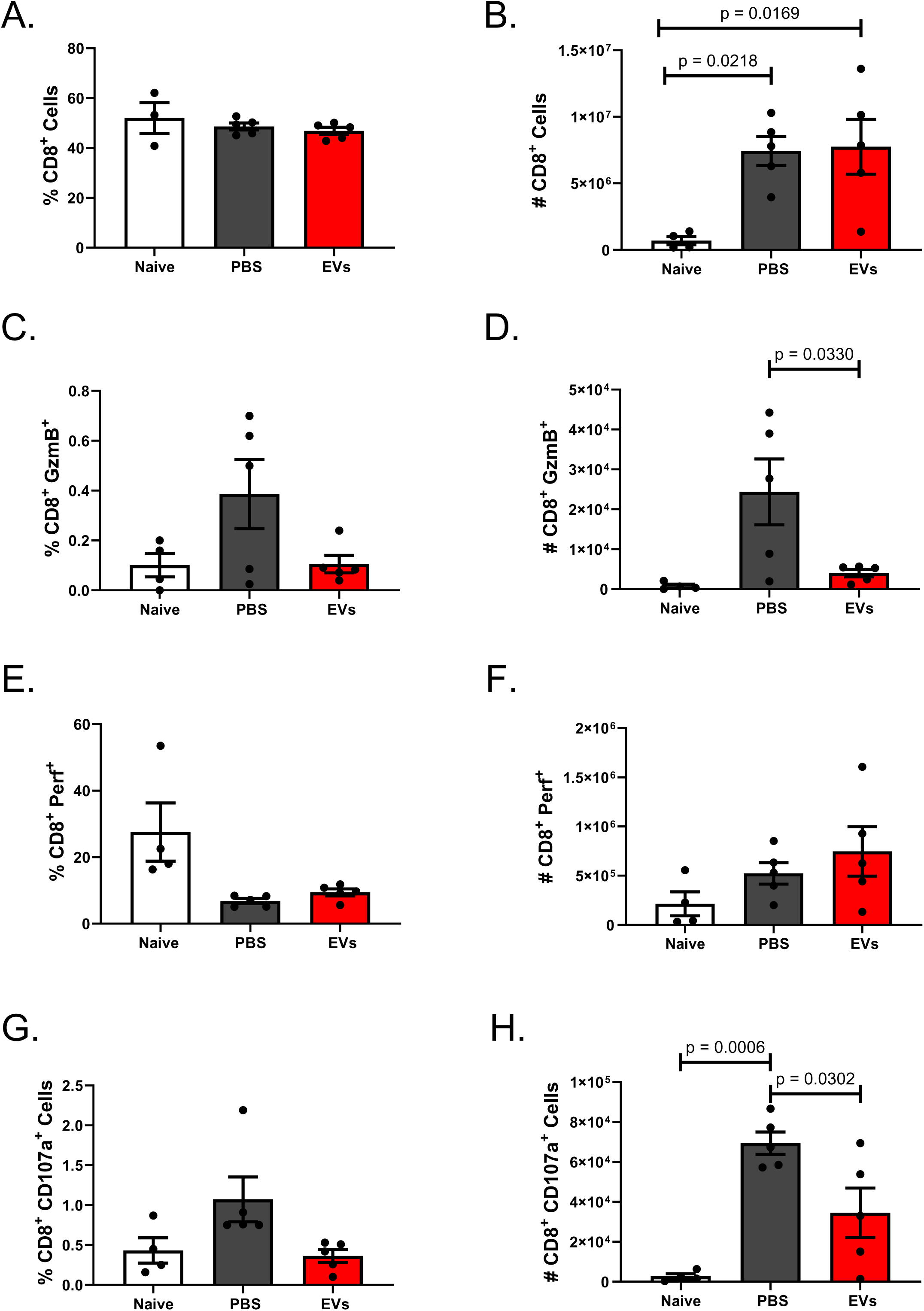
CD8⁺ T-cell cytotoxic responses following AD-MSC-EVs therapy. Draining lymph nodes were analyzed on day 43. (A, B) Frequency and number of CD8⁺ T cells; (C, D) Granzyme⁺CD8⁺ cells; (E, F) Perforin⁺CD8⁺ cells; (G, H) CD107a⁺CD8⁺ cells. n=4 (naïve), n=5 (PBS, EVs). Only p < 0.05 is shown. One-way ANOVA with Tukey’s post-test.

### Combined therapy with AD-MSC-EVs and pentavalent antimonial enhances lesion control without improving parasite clearance

To determine whether EVs could potentiate antimonial therapy, mice received intralesional EVs and/or meglumine antimoniate (PA) beginning on day 18 post-infection. EV treatment alone reduced lesion size consistent with prior experiments. PA alone also reduced lesion size relative to PBS control. Mice receiving combined therapy demonstrated the greatest reduction in lesion size (Figure 8A). AUC analysis confirmed enhanced disease control relative to individual treatments (Figure S6). Representative images corroborated these findings (Figure 8B). PA monotherapy significantly reduced parasite load in the infected footpad but not in the draining lymph node (Figure 8C–D). The combined regimen did not further reduce parasite load in any tissue examined (Figure 8C–E) indicating that under this protocol, the use of EVs abrogated the antiparasitic effect of PA.

### Cytokine levels in footpad tissues reflect distinct modulatory effects of EVs and combined therapy

Cytokine concentrations in footpad supernatants revealed distinct patterns across treatment groups. Combined therapy increased IL-4 levels, an effect not observed with either therapy alone (Figure 9A). EV treatment elevated IL-10, and the combination with PA resulted in the highest IL-10 concentrations (Figure 9B). IL-17 levels were unchanged by any treatment (Figure 9C). IL-22 increased only in the combined treatment group (Figure 9D). IFN-γ was significantly increased by PA, with further elevation when PA was combined with EVs (Figure 9E). Collectively, these findings indicate a complex immunoregulatory environment within lesion homogenates, characterized by the concurrent accumulation of pro- and anti-inflammatory cytokines that contribute to improved lesion control despite ineffective parasite clearance.

### AD-MSC-EV–mediated control of lesion progression requires IL-10

To assess whether IL-10 is required for the therapeutic effect of AD-MSC-EVs, WT and IL-10–deficient mice were treated by intralesional administrations following the same regimen as WT mice. EV treatment reduced lesion size in WT mice, consistent with previous results (Figure 10A). In IL-10–/– mice, EV treatment failed to confer protection and was associated with larger lesion size at later time points (Figure 10B). Representative images illustrate these differences (Figure 10C). Parasite load in footpad, spleen, and draining lymph nodes did not differ between EV-treated and control groups in either genotype (Figure 10D–F). These data demonstrate that the protective effect of AD-MSC-EVs depends on endogenous production of the cytokine IL-10.

**Figure 10.**
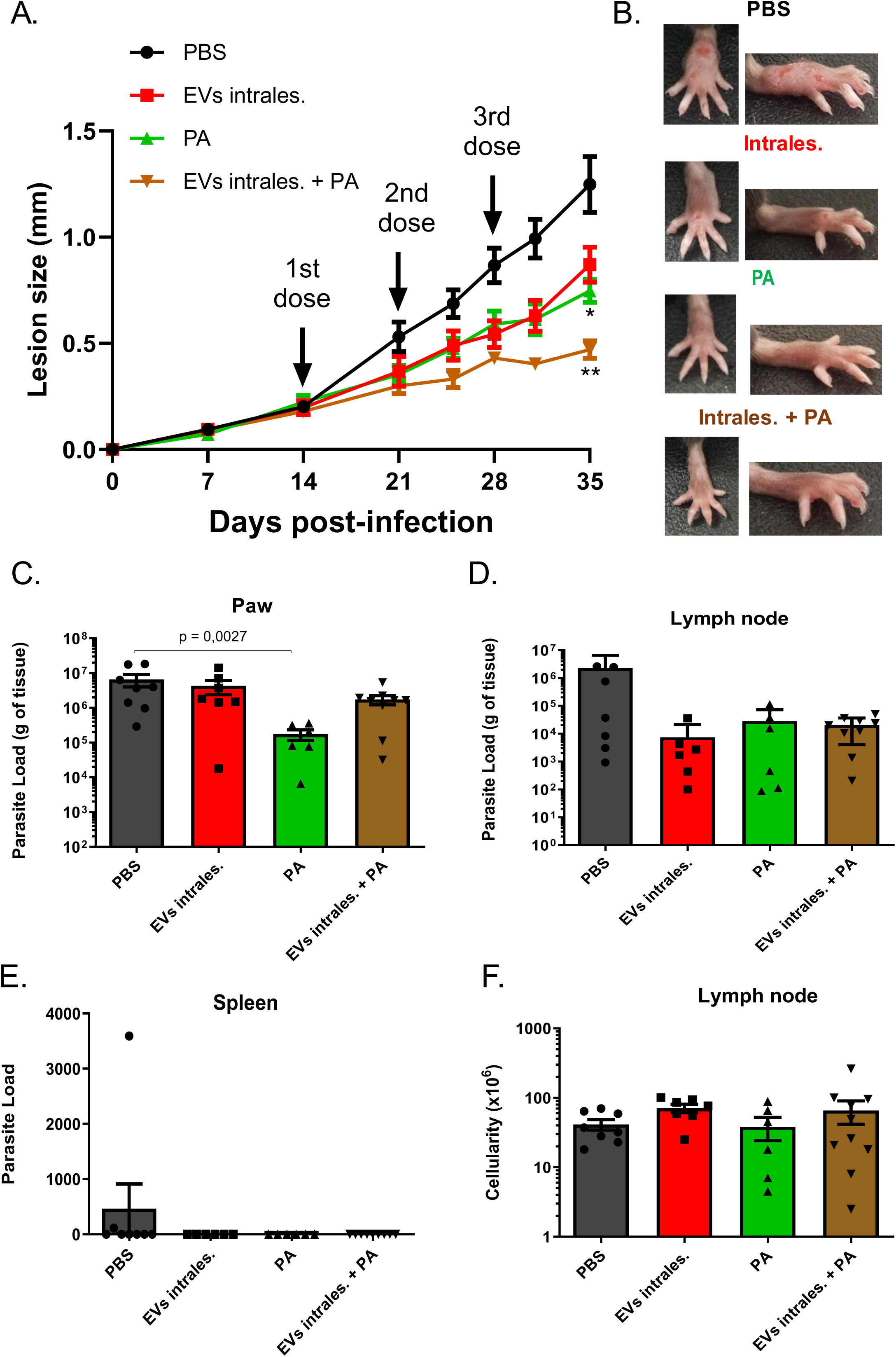
Combined AD-MSC-EVs and pentavalent antimonial therapy in *L. amazonensis*-infected mice. EVs (10⁹) were administered intralesionally (days 14, 21, 28). Pentavalent antimonial began on day 18. (A) Lesion progression; (B) Representative paw images; (C–E) Parasite load in paw, lymph nodes, and spleen; (F) Total lymph node cell count. Only p < 0.05 is shown.

**Figure 11.**
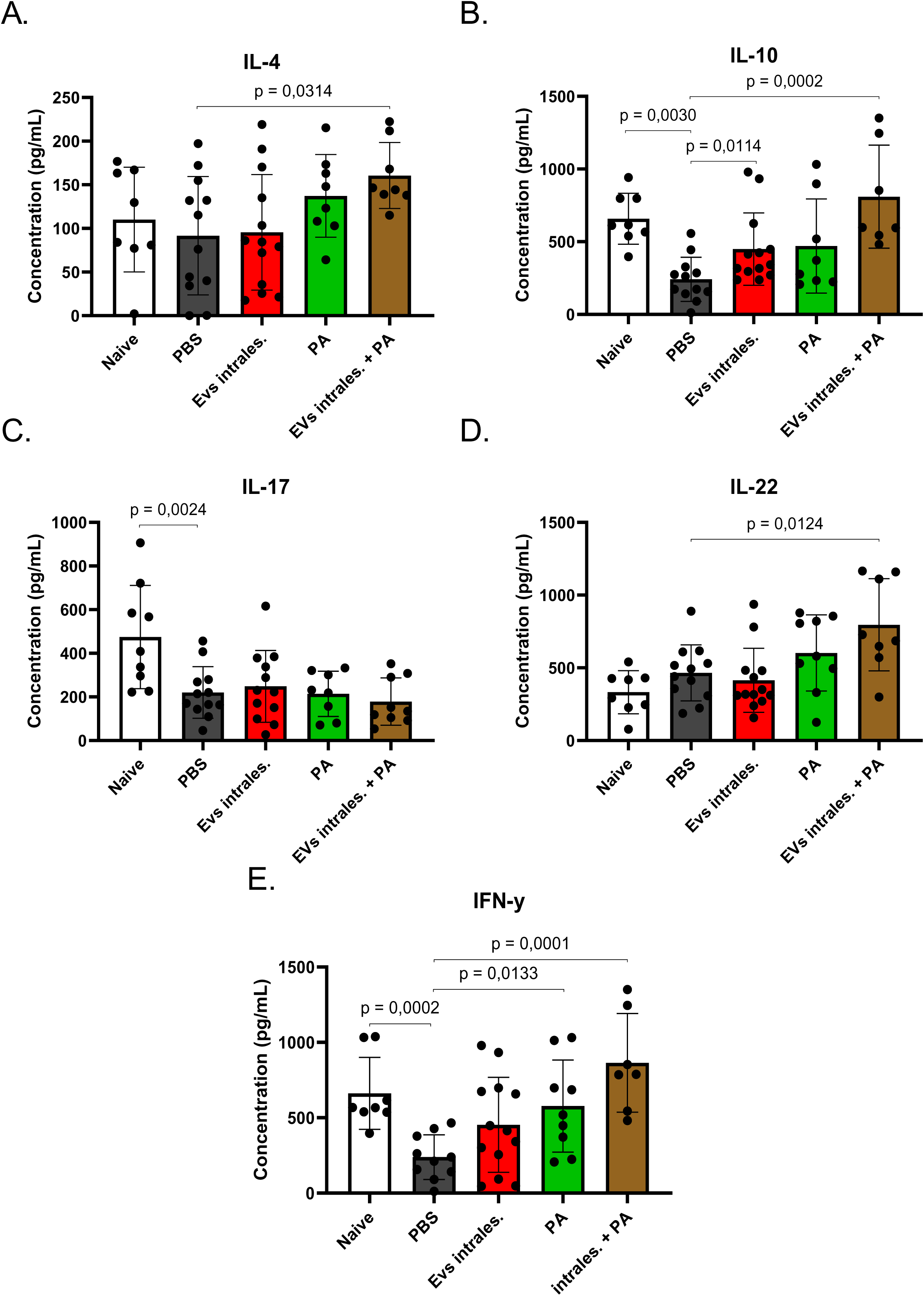
Cytokine levels at the lesion site. Cytokines in paw homogenates were measured following EV and/or antimonial treatment. (A) IL-4; (B) IL-10; (C) IL-17; (D) IL-22; (E) IFN-γ. Only p < 0.05 is shown (Mann–Whitney test).

**Figure 12.**
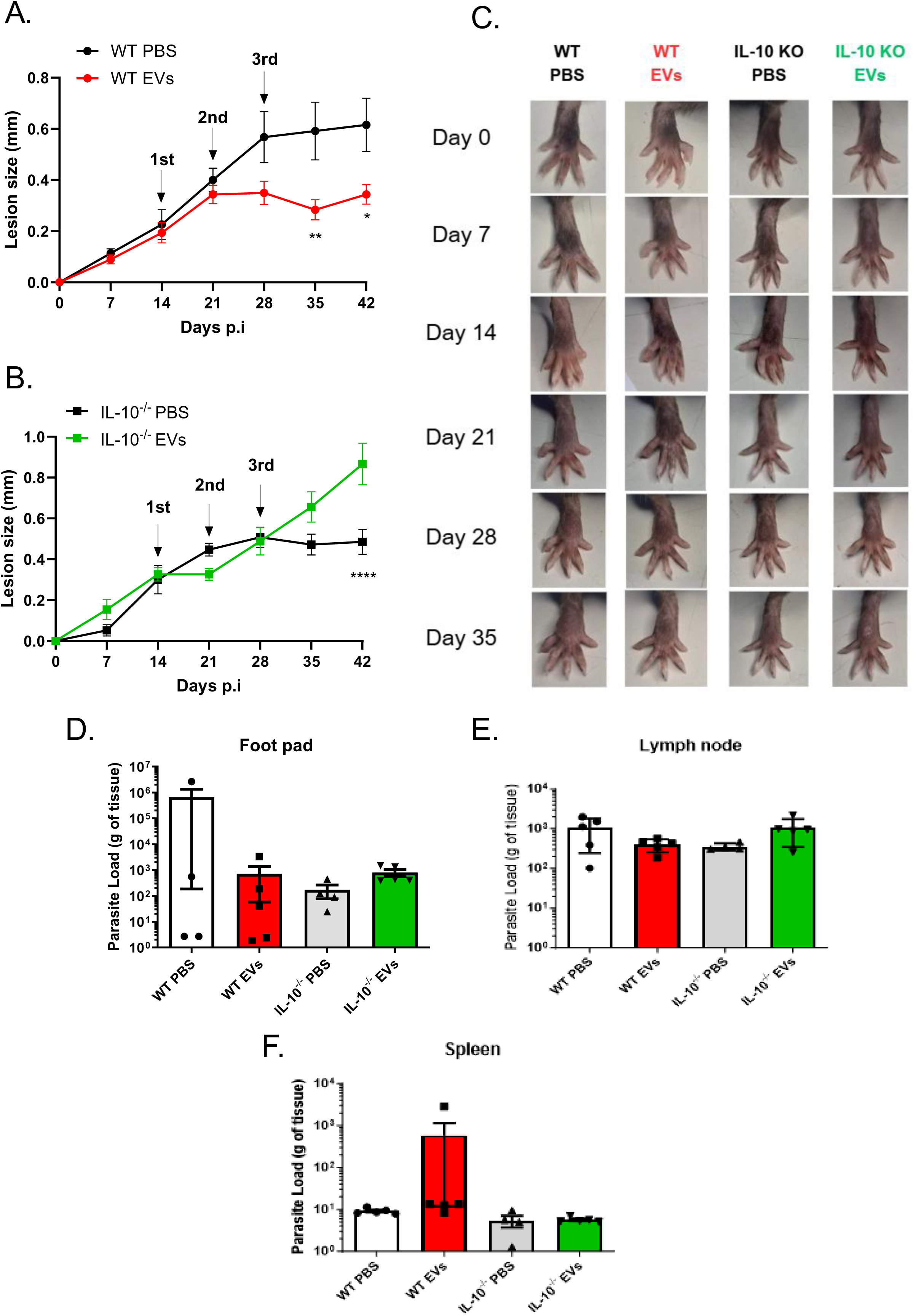
AD-MSC-EVs therapeutic effects require endogenous IL-10. WT and IL-10⁻/⁻ mice infected with *L. amazonensis* received EVs (10⁹) intralesionally. (A, B) Lesion progression in WT and IL-10⁻/⁻ mice; (C) Representative images; (D–F) Parasite load in footpad, lymph nodes, and spleen at day 43. Data are mean ± SEM from one of two experiments. *p < 0.01, **p < 0.001, ***p < 0.0001 vs. PBS; #p < 0.01 vs. naïve. Two-way ANOVA with Bonferroni post-test.

## Discussion

This study evaluated the therapeutic activity of extracellular vesicles derived from adipose-derived mesenchymal stem cells (AD-MSC-EVs) in *Leishmania amazonensis* infection, with emphasis on early parasite–macrophage interactions, modulation of host immunity, and the role of IL-10. The rationale for investigating these vesicles is supported by a growing body of evidence demonstrating that MSC-EVs reproduce the anti-inflammatory and reparative properties of their parent cells while offering practical advantages for clinical translation (Comariţa IK et al., 2022). In diverse inflammatory and infectious conditions, MSC-EVs suppress pro-inflammatory cytokines, limit immune-cell infiltration, regulate oxidative stress pathways, and favor M2-type macrophage polarization (HARRELL et al., 2019). These effects are particularly relevant in cutaneous leishmaniasis, in which persistent inflammation, dysregulated macrophage responses, and IL-10–dependent pathways contribute to tissue damage.

EVs isolated in this study fulfilled accepted criteria for size, morphology, and expression of canonical tetraspanins, thereby ensuring that subsequent *in vitro* and *in vivo* findings could be interpreted with confidence. In macrophages infected with *L. amazonensis*, AD-MSC-EVs reduced parasite load only when administered early after infection. This effect was independent of nitric oxide and instead associated with increased reactive oxygen species, indicating that EVs act within a narrow temporal window during the establishment of infection. Such early activity may influence vacuole formation or parasite differentiation. These observations parallel mechanisms described for antimonial agents, which rely on early ROS-dependent pathways (Mookerjee Basu J et al., 2006). Whether if specific EV cargos enhance ROS production or deliver leishmanicidal factors remains to be determined.

Pre-stimulation of macrophages with IFN-γ diminished the efficacy of EVs, despite enhanced NO generation. IFN-γ is known to accelerate parasitophorous vacuole maturation, a process that can favor intracellular parasite survival (Qi H, Ji J, Wanasen N, Soong L., 2004). The reduced sensitivity to EVs under these conditions highlights the importance of macrophage activation state and mirrors earlier studies in which AD-MSCs skewed macrophages toward an IL-10–rich regulatory phenotype rather than a classically activated, NO-dependent profile (Khosrowpour et al., 2017).

The route of administration markedly influenced therapeutic outcomes. Retro-orbital intravenous delivery did not affect lesion size or parasite burden, consistent with the well-established biodistribution of EVs to highly perfused organs such as the liver, spleen, and lungs (Aimaletdinov & Gomzikova, 2022), and likely reflecting limited EV accumulation at the lesion site. In contrast, intralesional administration significantly reduced lesion progression while leaving parasite load unchanged. This dissociation between tissue pathology and parasite burden is consistent with our previous findings using whole MSCs (Ramos et al., 2019) and supports the concept that EVs act as key mediators of MSC-driven immunomodulation rather than as direct leishmanicidal agents.

AD-MSC-EVs also modulated humoral immune responses, leading to reduced levels of parasite-specific IgM and IgG. Antibody production is an active contributor to pathology in *L. amazonensis* infection (Wanasen N et al., 2008; Firmino-Cruz et al., 2018), and previous studies have shown that AD-MSCs limit plasmablast generation while promoting IL-10–producing regulatory B cells (Franquesa et al., 2015). The present findings suggest that EVs recapitulate, at least in part, this regulatory effect on B-cell compartments.

Analysis of T-cell responses further clarified the mechanisms underlying lesion improvement. AD-MSC-EV treatment reduced IFN-γ- and IL-17–producing CD4⁺ T cells, consistent with attenuation of Th1 and Th17 responses—both implicated in tissue damage in *L. amazonensis* infection, and similar finds were observed in the treatment of diseases such as lupus (Li C, Wu F, Mao J, et al., 2024). Reduced IFN-γ likely contributes to the unchanged parasite load, whereas reduced IL-17 aligns with smaller lesion size as demonstrated before by our group (Dos-Santos JS et al, 2025). The increase in IL-10–producing γδ T cells suggests induction of a regulatory network capable of restraining excessive cytotoxicity. γδ T-cell–derived IL-10 has been shown to limit pathogenic CD8⁺ T-cell activity in other infections such as listeriosis (Rhodes KA et al, 2008), a mechanism that may also operate here.

Cytokine analysis of lesion homogenate supernatants showed that EV treatment locally induced IL-10, whereas meglumine antimoniate promoted IFN-γ production. The combined treatment elicited a broader cytokine response, characterized by increased levels of IL-4, IL-10, IL-22, and IFN-γ. A mixed cytokine profile has been associated with poor control of parasite burden (Ji et al., 2002). Notably, the combined therapy improved lesion control without reducing parasite load, a pattern that parallels findings from MSC–antimonial combination therapy and supports the potential of pairing immunomodulatory and antiparasitic strategies. This phenomenon contrasts with our previous observations using combined treatment with PA and AD-MSCs (Ramos et al., 2019). This discrepancy may be explained by the use of whole cells *in vivo*, which can induce additional mediators that are absent in EV preparations.

The central role of IL-10 was confirmed using IL-10–deficient mice, in which AD-MSC-EVs failed to limit lesion development and, at later stages of infection, even exacerbated tissue damage. These findings establish IL-10 as essential for the therapeutic effect and strongly suggest that EV cargo mediates IL-10 induction in recipient cells. Identifying the specific EV-associated molecules responsible—whether proteins, mRNAs, or microRNAs—represents an important direction for future research.

### Strengths and Limitations

Strengths of this study include rigorous EV characterization aligned with ISEV guidelines, systematic integration of mechanistic and in vivo analyses, and the use of IL-10–deficient mice to delineate cytokine dependence. The comparison of intralesional and intravenous routes also provides relevant translational information for future therapeutic development. Limitations include the absence of EV biodistribution studies following intralesional delivery, the lack of longitudinal parasite quantification, and incomplete identification of the EV cargo responsible for IL-10 induction. Moreover, while the C57BL/6 *L. amazonensis* model captures key features of disease pathogenesis, its translation to human leishmaniasis requires consideration of differences in host immunity and chronic disease kinetics.

## Conclusion

AD-MSC-EVs mitigate tissue pathology in *L. amazonensis* infection through IL-10–dependent immunomodulation, without significantly altering parasite load. Their ability to reduce lesion severity and to potentiate the benefits of meglumine antimoniate in the lesion control underscores their potential as adjunctive therapy in cutaneous leishmaniasis. Further studies focusing on EV cargo, optimized delivery strategies, and performance in chronic or treatment-resistant models will be essential for advancing this approach toward clinical application.

## Acknowledgments

We would like to thank the facilities at the Centro Nacional de Biologia Estrutural e Bioimagem (CENABIO) and the Unidade de Microscopia Multiusuário Padrón Lins IMPG (Unimicro) for their assistance with sample preparation and TEM analysis. Additionally, we acknowledge the support of the Instituto de Biofísica Carlos Chagas Filho (IBCCF) and the Instituto de Microbiologia Paulo de Goés for their contributions to flow cytometry and NTA acquisitions.

## Funding

This study was supported by the Oswaldo Cruz Institute (IOC - Fiocruz), Brazilian Council for Scientific and Technological Development (CNPq), the Rio de Janeiro State Research Foundation (FAPERJ) and, the Coordination for the Improvement of Higher Education Personnel (CAPES).

## Author contributions

Conception and design, DBA, HLMG, TDR; Collection and/or assembly of data, AAR, IBS, JVPR, HPS, NCSM, RTS, BTM; Data analysis and interpretation, DBA, RTS, TDR, BTM, LC, DG, FFC, AMFM; Manuscript writing, DBA, TDR, HLMG, FFC, PR; Provision of study material, AMFM, FFC, HLMG, DG, LC, PR; Final approval of manuscript, TDR, FFC, PR, HLMG; Administrative support, JSS; Financial support, AMFM, PR, HLMG. All authors read and approved the final manuscript.

## Data availability

All data needed to evaluate the conclusions in the manuscript are present in the manuscript and the supplementary Materials. All methods and protocols used are included in the main manuscript. All reagents, antibodies, and resources are listed in the main manuscript or supplementary files. The data sets used and/or analyzed during the current study are available from the corresponding authors on reasonable request.

## Ethical approval and consent to participate

This prospective, randomized experimental study was approved by the Local Animal Research Ethics Committee (CEUA CCS-110/17, 2017) of the Health Sciences Centre of the Federal University of Rio de Janeiro (UFRJ), Rio de Janeiro, Brazil. All animals received care in accordance with the “Principles of Laboratory Animal Care” formulated by the National Society for Medical Research (Bethesda, Maryland) and the U.S. National Academy of Sciences (Bethesda, Maryland) Guide for the Care and Use of Laboratory Animals. This study followed the Animal Research: Reporting of In Vivo Experiments (ARRIVE) guidelines for animal experimentation (Percie du Sert N *et al*., 2020) Mice were housed at a controlled temperature (23°C) and controlled light–dark cycle (12–12 h), with free access to water and food; no acclimation was done.

## Consent for publication

Not applicable.

## Competing interests

The authors declare no competing interest.

## Supplementary Figures Legends

Figure S1 - ***In vitro* effects of MSC EVs on infected macrophages.** Macrophages were infected for 4 h and treated with EVs for 24 or 48 h. (A, D) Percentage of infected macrophages; (B, E) Internalized amastigotes; (C, F) Nitrite levels. Data are mean ± SEM from two experiments. *p < 0.01, **p < 0.001, ***p < 0.0001. One-way ANOVA with Tukey’s post-test.

Figure S2. **MSC EVs in IFN-γ–stimulated, *L. amazonensis*-infected macrophages.** Macrophages were infected for 4 h, stimulated with IFN-γ at hour 3, and treated with EVs at hour 4. (A, D) Percentage of infected macrophages; (B, E) Internalized amastigotes; (C, F) Nitric oxide levels. CTR NI+10⁹: non-infected macrophages treated with EVs; CTR NI+IFN: non-infected macrophages stimulated with IFN-γ; NT+IFN: infected macrophages stimulated with IFN-γ.

Figure S3. **Gating strategy for flow cytometric analyses.** Draining lymph nodes were collected from infected mice at euthanasia, mechanically dissociated, filtered, and stained as described in the Methods. The scheme illustrates the gating strategy used to identify CD4⁺ and CD8⁺ T-cell populations and to quantify cytokine-producing subsets following treatment.

Figure S4. **CD8⁺ T-cell cytokine responses following intralesional administration of MSC extracellular vesicles in *L. amazonensis*–infected mice.** C57BL/6 mice were infected in the right hind-paw footpad with 2×10⁶ *L. amazonensis* promastigotes and received three intralesional doses of MSC-derived EVs (1×10⁹) on days 14, 21, and 28 post-infection. The control group received PBS. On day 43, draining lymph nodes from naïve, PBS-treated, and EV-treated mice were harvested, mechanically dissociated, and stained for flow cytometry. (A, B) Frequency and number of CD8⁺ T cells gated on CD3⁺ cells; (C, D) IFN-γ⁺CD8⁺ cells; (E, F) IL-17⁺CD8⁺ cells; (G, H) IL-10⁺CD8⁺ cells. Naïve (n=4), PBS (n=5), EV-treated (n=5). Only statistically significant differences (p < 0.05) are indicated. One-way ANOVA with Tukey’s post-test (GraphPad Prism).

Figure S5. **Area under the curve (AUC) of lesion progression in mice receiving EVs alone or in combination with pentavalent antimonial.** C57BL/6 mice were infected in the right hind-paw footpad with 2×10⁶ *L. amazonensis* promastigotes. EV-treated groups received three intralesional doses of MSC-derived EVs (1×10⁹) on days 14, 21, and 28 post-infection. Mice assigned to pentavalent antimonial (PA) therapy received intraperitoneal doses on alternate days beginning on day 18. Lesion development was monitored by pachymetry, and the area under the curve (AUC) was calculated to compare overall lesion burden across treatment groups.

## Notes

### Competing Interest Statement

The authors have declared no competing interest.

## References

Aimaletdinov AM, Gomzikova MO. Tracking of Extracellular Vesicles’ Biodistribution: New Methods and Approaches. Int J Mol Sci. 2022;23(19):11312. Published 2022 Sep 25. doi:10.3390/ijms231911312

Bahrami S, Safari M, Razi Jalali MH, Ghorbanpoor M, Tabandeh MR, Rezaie A. The potential therapeutic effect of adipose-derived mesenchymal stem cells in the treatment of cutaneous leishmaniasis caused by L. major in BALB/c mice. Exp Parasitol. 2021;222:108063. doi:10.1016/j.exppara.2020.108063

Mookerjee Basu J, Mookerjee A, Sen P, Bhaumik S, Sen P, Banerjee S, Naskar K, Choudhuri SK, Saha B, Raha S, Roy S. Sodium antimony gluconate induces generation of reactive oxygen species and nitric oxide via phosphoinositide 3-kinase and mitogen-activated protein kinase activation in Leishmania donovani-infected macrophages. Antimicrob Agents Chemother. 2006 May;50(5):1788–97. doi: 10.1128/AAC.50.5.1788-1797.2006. PMID: 16641451; PMCID: PMC1472228.

Bharadava K, Upadhyay TK, Kaushal RS, et al. Genomic Insight of Leishmania Parasite: In-Depth Review of Drug Resistance Mechanisms and Genetic Mutations. ACS Omega. 2024;9(11):12500–12514. Published 2024 Mar 8. doi:10.1021/acsomega.3c09400

Cho BS, Kim JO, Ha DH, Yi YW. Exosomes derived from human adipose tissue-derived mesenchymal stem cells alleviate atopic dermatitis. Stem Cell Res Ther. 2018;9:187. doi: 10.1186/s13287-018-0939-5.

Comariţa IK, Vîlcu A, Constantin A, Procopciuc A, Safciuc F, Alexandru N, et al. Therapeutic potential of stem cell-derived extracellular vesicles on atherosclerosis-induced vascular dysfunction and its key molecular players. Front Cell Dev Biol. 2022;10:817180. doi: 10.3389/fcell.2022.817180.

Dos-Santos JS, Firmino-Cruz L, Oliveira-Maciel D, et al. IL-17A/IFN-γ producing γδ T cell functional dichotomy impacts cutaneous leishmaniasis in mice. J Leukoc Biol. 2025;117(3):qiae251. doi:10.1093/jleuko/qiae251

Firmino-Cruz L, Ramos TD, da Fonseca-Martins AM, et al. Immunomodulating role of IL-10-producing B cells in Leishmania amazonensis infection. Cell Immunol. 2018;334:20–30. doi:10.1016/j.cellimm.2018.08.014

Fischer T, Fischer M, Schliemann S, Elsner P. Treatment of mucocutaneous leishmaniasis - A systematic review. J Dtsch Dermatol Ges. Published online May 20, 2024. doi:10.1111/ddg.15424

Franquesa M, Mensah FK, Huizinga R, Strini T, Boon L, Lombardo E, DelaRosa O, Laman JD, Grinyó JM, Weimar W, Betjes MG, Baan CC, Hoogduijn MJ. Human adipose tissue-derived mesenchymal stem cells abrogate plasmablast formation and induce regulatory B cells independently of T helper cells. Stem Cells. 2015 Mar;33(3):880–91. doi: 10.1002/stem.1881. PMID: 25376628.

Habiba UE, Khan N, Greene DL, Ahmad K, Shamim S, Umer A. Meta-analysis shows that mesenchymal stem cell therapy can be a possible treatment for diabetes. Front Endocrinol (Lausanne). 2024;15:1380443. Published 2024 May 10. doi:10.3389/fendo.2024.1380443

Harrell CR, Fellabaum C, Jovicic N, Djonov V, Arsenijevic N, Volarevic V. Molecular Mechanisms Responsible for Therapeutic Potential of Mesenchymal Stem Cell-Derived Secretome. Cells. 2019;8(5):467. Published 2019 May 16. doi:10.3390/cells8050467

Harrell C.R., Jovicic N., Djonov V., Arsenijevic N., Volarevic V. Mesenchymal stem cell-derived exosomes and other extracellular vesicles as new remedies in the therapy of inflammatory diseases. Cells. 2019;8

Khosrowpour Z, Hashemi SM, Mohammadi-Yeganeh S, Soudi S. Pretreatment of Mesenchymal Stem Cells With Leishmania major Soluble Antigens Induce Anti-Inflammatory Properties in Mouse Peritoneal Macrophages. J Cell Biochem. 2017;118(9):2764–2779. doi:10.1002/jcb.25926

Li C, Wu F, Mao J, et al. Mesenchymal stem cells-derived extracellular vesicles ameliorate lupus nephritis by regulating T and B cell responses. Stem Cell Res Ther. 2024;15(1):216. Published 2024 Jul 18. doi:10.1186/s13287-024-03834-w

Li Y, Altemus J, Lightner AL. Mesenchymal stem cells and acellular products attenuate murine induced colitis. Stem Cell Res Ther. 2020;11(1):515. Published 2020 Nov 30. doi:10.1186/s13287-020-02025-7

Maspi N, Abdoli A, Ghaffarifar F. Pro- and anti-inflammatory cytokines in cutaneous leishmaniasis: a review. Pathog Glob Health. 2016;110(6):247–260. doi:10.1080/20477724.2016.1232042

Mello DB, Ramos IP, Mesquita FC, Brasil GV, Rocha NN, Takiya CM, et al. Adipose tissue-derived Mesenchymal stromal cells protect mice infected with Trypanosoma cruzi from cardiac damage through modulation of anti-parasite immunity. PLoS Negl Trop Dis. 2015;9:e0003945.

Percie du Sert N, Hurst V, Ahluwalia A, Alam S, Avey MT, Baker M, Browne WJ, Clark A, Cuthill IC, Dirnagl U, Emerson M, Garner P, Holgate ST, Howells DW, Karp NA, Lazic SE, Lidster K, MacCallum CJ, Macleod M, Pearl EJ, Petersen OH, Rawle F, Reynolds P, Rooney K, Sena ES, Silberberg SD, Steckler T, Würbel H. The ARRIVE guidelines 2.0: Updated guidelines for reporting animal research. PLoS Biol. 2020 Jul 14;18(7):e3000410. doi: 10.1371/journal.pbio.3000410. PMID: 32663219; PMCID: PMC7360023.

Pereira JC, Ramos TD, Silva JD, et al. Effects of Bone Marrow Mesenchymal Stromal Cell Therapy in Experimental Cutaneous Leishmaniasis in BALB/c Mice Induced by *Leishmania amazonensis*. Front Immunol. 2017;8:893. Published 2017 Aug 10. doi:10.3389/fimmu.2017.00893

Pinheiro AAS, Torrecilhas AC, Souza BSF, Cruz FF, Guedes HLM, Ramos TD, Lopes-Pacheco M, Caruso-Neves C, Rocco PRM. Potential of extracellular vesicles in the pathogenesis, diagnosis and therapy for parasitic diseases. J Extracell Vesicles. 2024 Aug;13(8):e12496. doi: 10.1002/jev2.12496. PMID: 39113589; PMCID: PMC11306921.

Qi H, Ji J, Wanasen N, Soong L. Enhanced replication of Leishmania amazonensis amastigotes in gamma interferon-stimulated murine macrophages: implications for the pathogenesis of cutaneous leishmaniasis. Infect Immun. 2004 Feb;72(2):988–95. doi: 10.1128/IAI.72.2.988-995.2004. PMID: 14742545; PMCID: PMC321581.

Quan J, Liu Q, Li P, Yang Z, Zhang Y, Zhao F, Zhu G. Mesenchymal stem cell exosome therapy: current research status in the treatment of neurodegenerative diseases and the possibility of reversing normal brain aging. Stem Cell Res Ther. 2025 Feb 21;16(1):76. doi: 10.1186/s13287-025-04160-5. PMID: 39985030; PMCID: PMC11846194.

Ramos TD, Silva JD, da Fonseca-Martins AM, et al. Combined therapy with adipose tissue-derived mesenchymal stromal cells and meglumine antimoniate controls lesion development and parasite load in murine cutaneous leishmaniasis caused by Leishmania amazonensis. Stem Cell Res Ther. 2020;11(1):374. Published 2020 Aug 31. doi:10.1186/s13287-020-01889-z

Regmi S, Pathak S, Kim JO, Yong CS, Jeong JH. Mesenchymal stem cell therapy for the treatment of inflammatory diseases: Challenges, opportunities, and future perspectives. Eur J Cell Biol. 2019;98(5-8):151041. doi:10.1016/j.ejcb.2019.04.002

Rhodes KA, Andrew EM, Newton DJ, Tramonti D, Carding SR. A subset of IL-10-producing gammadelta T cells protect the liver from Listeria-elicited, CD8(+) T cell-mediated injury. Eur J Immunol. 2008;38(8):2274–2283. doi:10.1002/eji.200838354

Rodríguez-Serrato MA, Salinas-Carmona MC, Limón-Flores AY. Immune response to Leishmania mexicana: the host-parasite relationship. Pathog Dis. 2020;78(8):ftaa060. doi:10.1093/femspd/ftaa060

Serrano-Coll H, Aristizábal-Parra LK, Olarte G, Salamanca-Leguizamón C. Cutaneous leishmaniasis: immunological insights and clinical challenges. Colomb Med (Cali). 2025;56(3):e3006750. Published 2025 Sep 15. doi:10.25100/cm.v56i3.6750

Song Y, Liang F, Tian W, Rayhill E, Ye L, Tian X. Optimizing therapeutic outcomes: preconditioning strategies for MSC-derived extracellular vesicles. Front Pharmacol. 2025 Feb 10;16:1509418. doi: 10.3389/fphar.2025.1509418. PMID: 39995418; PMCID: PMC11847897.

Thakur RS, Awasthi V, Sanyal A, et al. Mesenchymal stem cells protect against malaria pathogenesis by reprogramming erythropoiesis in the bone marrow. Cell Death Discov. 2020;6(1):125. Published 2020 Nov 15. doi:10.1038/s41420-020-00363-2

Wanasen N, Xin L, Soong L. Pathogenic role of B cells and antibodies in murine Leishmania amazonensis infection. Int J Parasitol. 2008;38(3-4):417–429. doi:10.1016/j.ijpara.2007.08.010

Welsh JA, Goberdhan DCI, O’Driscoll L, Buzas EI, Blenkiron C, Bussolati B, Cai H, Di Vizio D, Driedonks TAP, Erdbrügger U, Falcon-Perez JM, Fu QL, Hill AF, Lenassi M, Lim SK, Mahoney MG, Mohanty S, Möller A, Nieuwland R, Ochiya T, Sahoo S, Torrecilhas AC, Zheng L, Zijlstra A, Abuelreich S, Bagabas R, Bergese P, Bridges EM, Brucale M, Burger D, Carney RP, Cocucci E, Crescitelli R, Hanser E, Harris AL, Haughey NJ, Hendrix A, Ivanov AR, Jovanovic-Talisman T, Kruh-Garcia NA, Ku’ulei-Lyn Faustino V, Kyburz D, Lässer C, Lennon KM, Lötvall J, Maddox AL, Martens-Uzunova ES, Mizenko RR, Newman LA, Ridolfi A, Rohde E, Rojalin T, Rowland A, Saftics A, Sandau US, Saugstad JA, Shekari F, Swift S, Ter-Ovanesyan D, Tosar JP, Useckaite Z, Valle F, Varga Z, van der Pol E, van Herwijnen MJC, Wauben MHM, Wehman AM, Williams S, Zendrini A, Zimmerman AJ; MISEV Consortium; Théry C, Witwer KW. Minimal information for studies of extracellular vesicles (MISEV2023): From basic to advanced approaches. J Extracell Vesicles. 2024 Feb;13(2):e12404. doi: 10.1002/jev2.12404. Erratum in: J Extracell Vesicles. 2024 May;13(5):e12451. doi: 10.1002/jev2.12451. PMID: 38326288; PMCID: PMC10850029.

Wei S, Li M, Wang Q, et al. Mesenchymal Stromal Cells: New Generation Treatment of Inflammatory Bowel Disease. J Inflamm Res. 2024;17:3307–3334. Published 2024 May 22. doi:10.2147/JIR.S458103

World Health Organization. Leishmaniasis. Who int; (2025). Available at: https://www.who.int/news-room/fact-sheets/detail/leishmaniasis

