## Supplemental figures for "Adipose-Derived Extracellular Vesicles Mitigate Experimental Cutaneous Leishmaniasis Through an IL-10–Dependent Mechanism"

### Figure S1

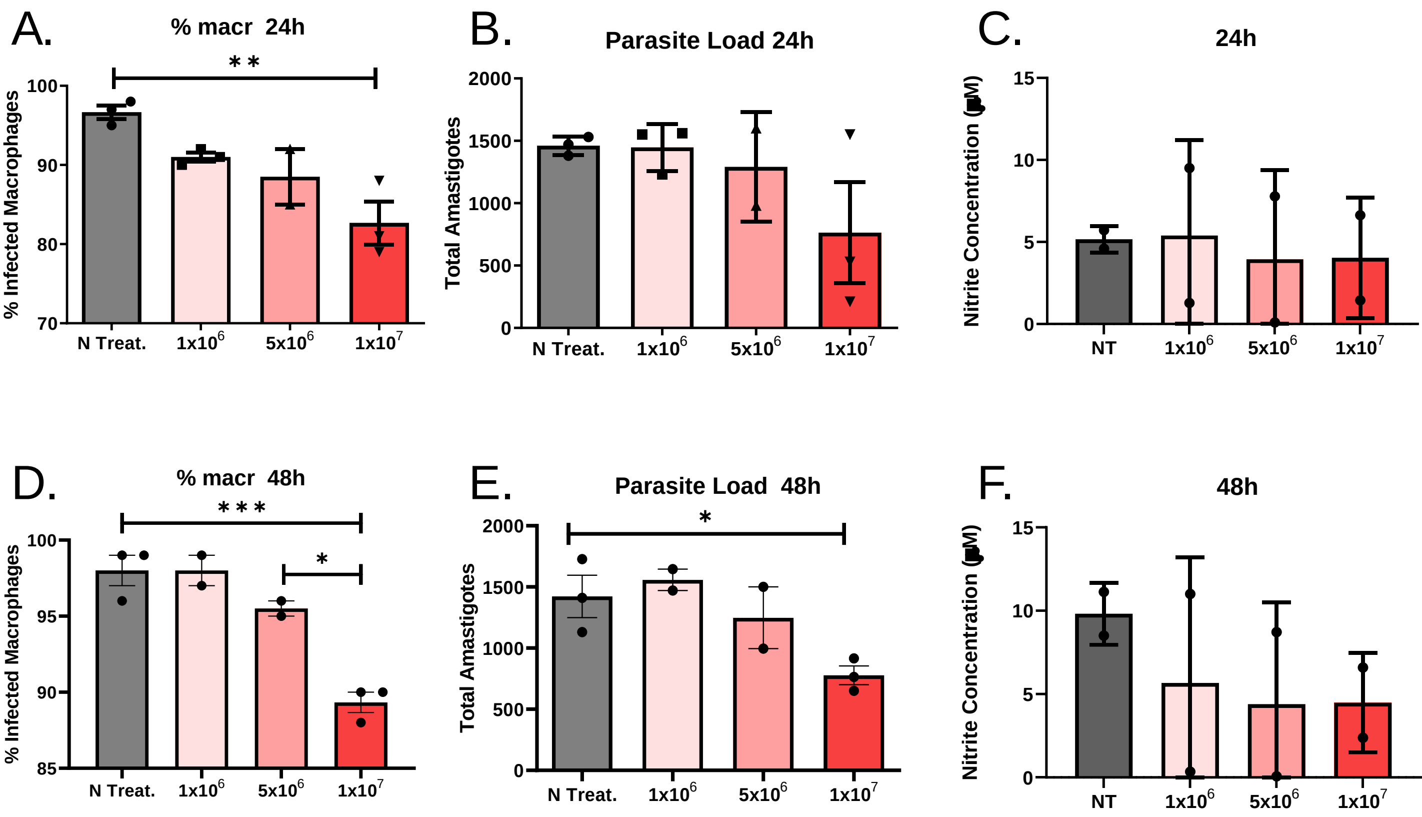
A.
% macr 24h
**
% Infected Macrophages
100
90
80
70
N Treat.
1x10^6
5x10^6
1x10^7
B.
Parasite Load 24h
Total Amastigotes
2000
1500
1000
500
0
N Treat.
1x10^6
5x10^6
1x10^7
C.
24h
Nitrite Concentration (μM)
15
10
5
0
NT
1x10^6
5x10^6
1x10^7
D.
% macr 48h
***
*
% Infected Macrophages
100
95
90
85
N Treat.
1x10^6
5x10^6
1x10^7
E.
Parasite Load 48h
*
Total Amastigotes
2000
1500
1000
500
0
N Treat.
1x10^6
5x10^6
1x10^7
F.
48h
Nitrite Concentration (μM)
15
10
5
0
NT
1x10^6
5x10^6
1x10^7

### Figure S2

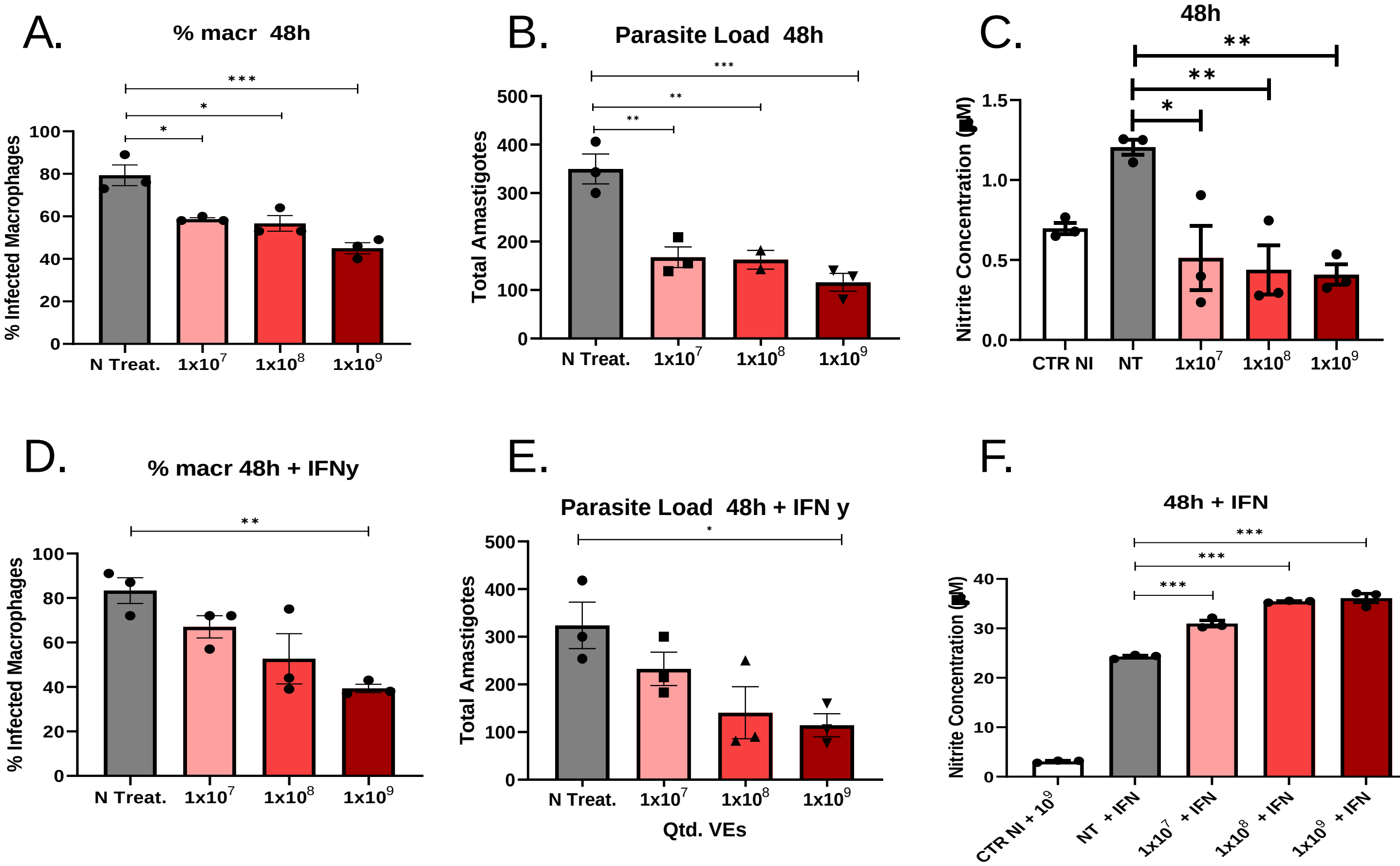

A.
% macr 48h
% Infected Macrophages
N Treat.
1x10^7
1x10^8
1x10^9
B.
Parasite Load 48h
Total Amastigotes
C.
48h
Nitrite Concentration (µM)
CTR NI
NT
D.
% macr 48h + IFNy
E.
Parasite Load 48h + IFN y
Qtd. VEs
F.
48h + IFN
CTR NI + 10^9
NT + IFN
1x10^7 + IFN
1x10^8 + IFN
1x10^9 + IFN

### Figure S3

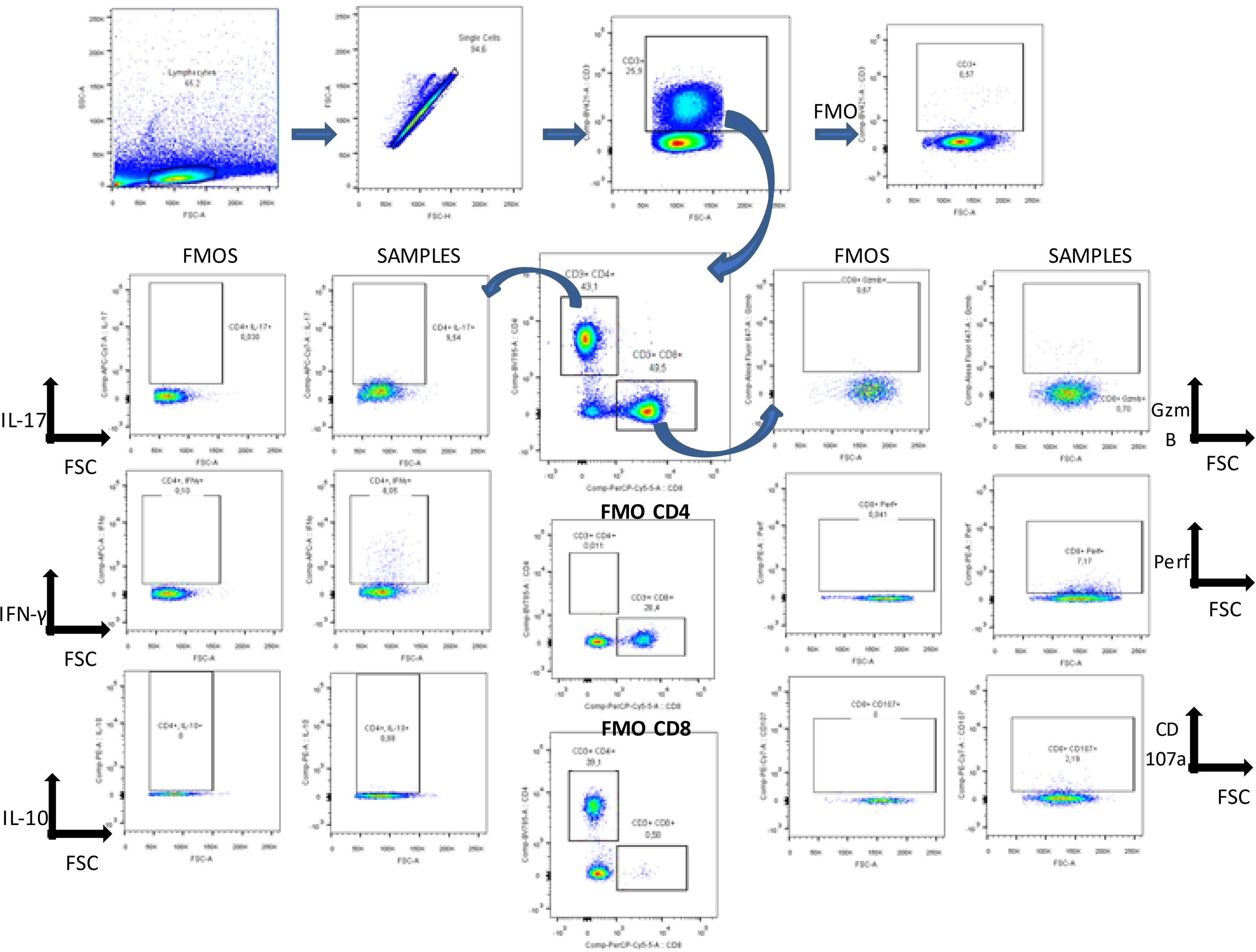


**Figure S4. Gate strategy for Flow Cytometry.** Draining lymph nodes of the lesion of infected mice were collected after euthanasia macerated and stained as described in the methods session. The scheme shows the analysis strategy used to observe CD4 and CD8 responses to treatment.

### Figure S4

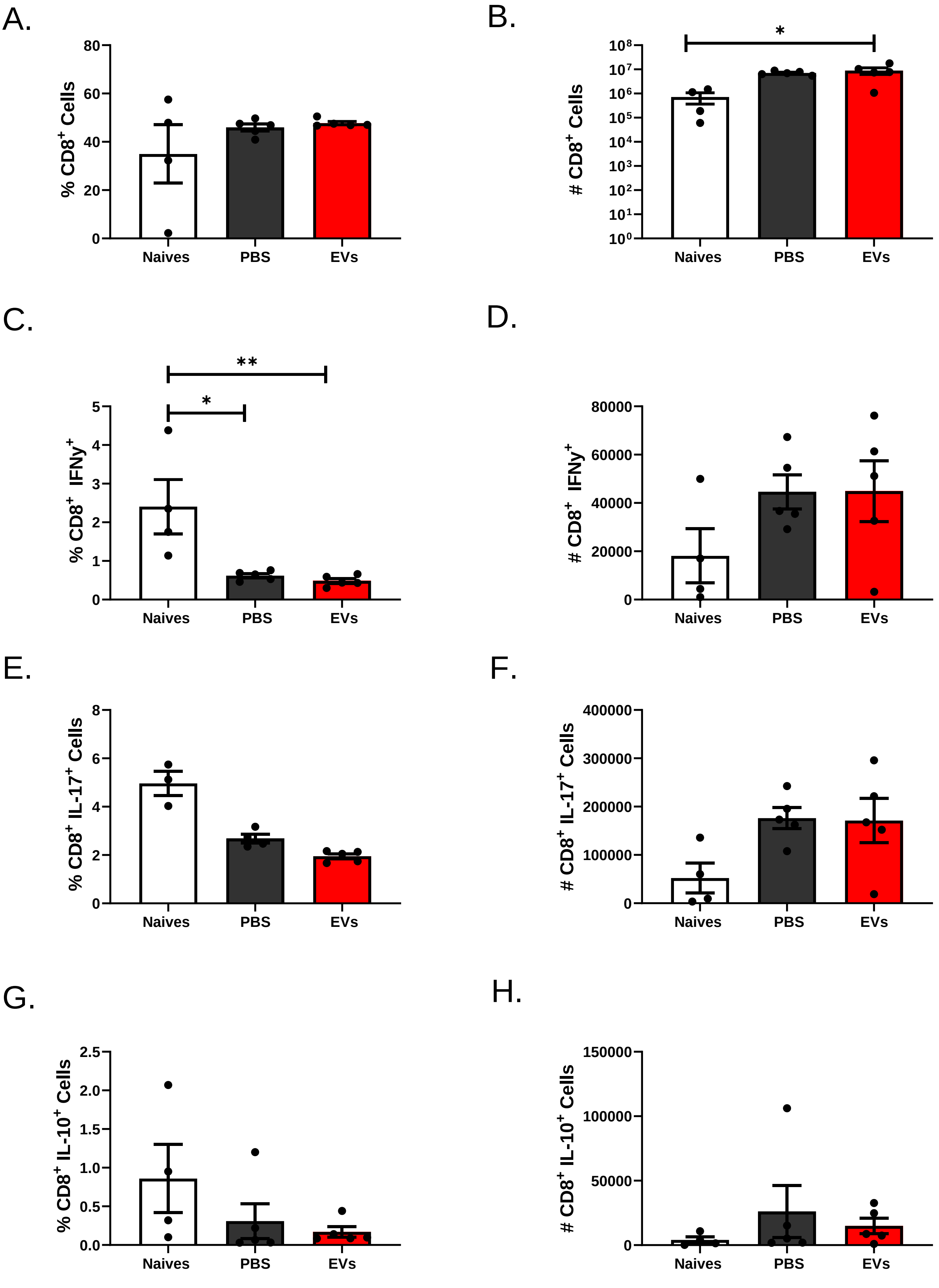

A.
% CD8+ Cells
0
20
40
60
80
Naives
PBS
EVs
B.
### CD8+ Cells
10⁰
10¹
10²
10³
10⁴
10⁵
10⁶
10⁷
10⁸
*
Naives
PBS
EVs
C.
% CD8+ IFNy+
0
1
2
3
4
5
**
*
Naives
PBS
EVs
D.
### CD8+ IFNy+
0
20000
40000
60000
80000
Naives
PBS
EVs
E.
% CD8+ IL-17+ Cells
0
2
4
6
8
Naives
PBS
EVs
F.
### CD8+ IL-17+ Cells
0
100000
200000
300000
400000
Naives
PBS
EVs
G.
% CD8+ IL-10+ Cells
0.0
0.5
1.0
1.5
2.0
2.5
Naives
PBS
EVs
H.
### CD8+ IL-10+ Cells
0
50000
100000
150000
Naives
PBS
EVs

**Figure S5**

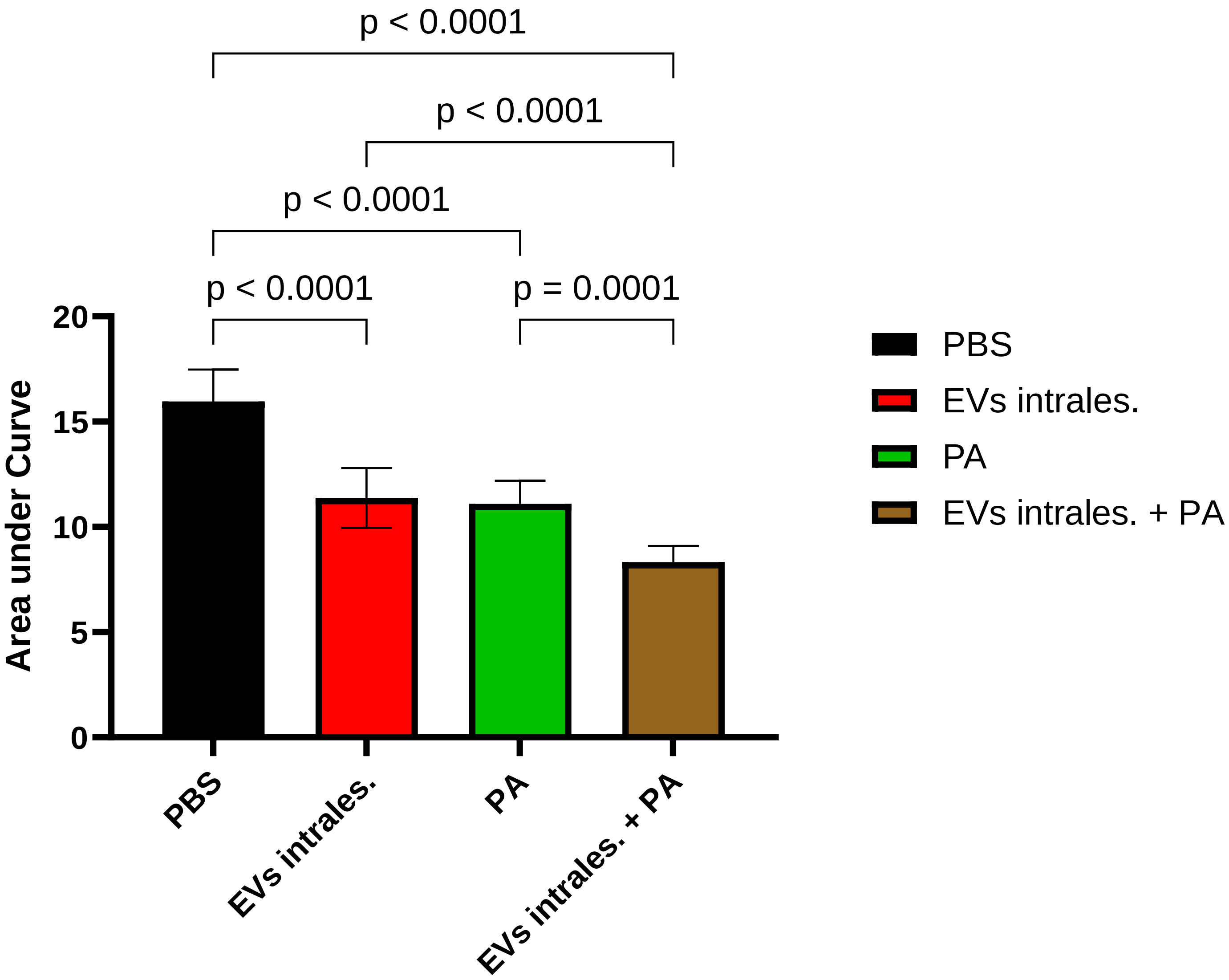

p < 0.0001
p < 0.0001
p < 0.0001
p < 0.0001
p = 0.0001
Area under Curve
20
15
10
5
0
PBS
EVs intrales.
PA
EVs intrales. + PA
PBS
EVs intrales.
PA
EVs intrales. + PA
